# Nonlinear manifolds underlie neural population activity during behaviour

**DOI:** 10.1101/2023.07.18.549575

**Authors:** Cátia Fortunato, Jorge Bennasar-Vázquez, Junchol Park, Joanna C. Chang, Lee E. Miller, Joshua T. Dudman, Matthew G. Perich, Juan A. Gallego

**Affiliations:** Department of Bioengineering, Imperial College London, London UK; Janelia Research Campus, Howard Hughes Medical Institute, Ashburn VA, USA; Department of Neurosciences, Northwestern University, Chicago IL, USA; Department of Biomedical Engineering, Northwestern University, Chicago IL, USA; Department of Physical Medicine and Rehabilitation, Northwestern University, Chicago IL, USA, and Shirley Ryan Ability Lab, Chicago, IL, USA; Department of Neurosciences, Faculté de médecine, Université de Montréal, Montréal, Québec, Canada; Québec Artificial Intelligence Institute (MILA), Montréal, Québec, Canada

## Abstract

There is rich variety in the activity of single neurons recorded during behaviour. Yet, these diverse single neuron responses can be well described by relatively few patterns of neural co-modulation. The study of such low-dimensional structure of neural population activity has provided important insights into how the brain generates behaviour. Virtually all of these studies have used linear dimensionality reduction techniques to estimate these population-wide co-modulation patterns, constraining them to a flat “neural manifold”. Here, we hypothesised that since neurons have nonlinear responses and make thousands of distributed and recurrent connections that likely amplify such nonlinearities, neural manifolds should be intrinsically nonlinear. Combining neural population recordings from monkey, mouse, and human motor cortex, and mouse striatum, we show that: 1) neural manifolds are intrinsically nonlinear; 2) their nonlinearity becomes more evident during complex tasks that require more varied activity patterns; and 3) manifold nonlinearity varies across architecturally distinct brain regions. Simulations using recurrent neural network models confirmed the proposed relationship between circuit connectivity and manifold nonlinearity, including the differences across architecturally distinct regions. Thus, neural manifolds underlying the generation of behaviour are inherently nonlinear, and properly accounting for such nonlinearities will be critical as neuroscientists move towards studying numerous brain regions involved in increasingly complex and naturalistic behaviours.

## Introduction

Behaviour is ultimately generated by the orchestrated activity of neural populations across the brain. An increasing number of studies show that the coordinated activity of tens or hundreds of neurons within a given brain region can be captured by relatively few covariation patterns, which we call *neural modes*^1, 2^. This observation holds strikingly well across a variety of species, brain regions, and tasks, from the locust olfactory system during odour presentation^3^, to the human frontal cortex during memory and categorization^4^. These low-dimensional activity patterns are thought to reflect fundamental constraints on neural population activity^1^ that arise due to biophysical phenomena including circuit connectivity^5–8^, neuromodulators^9, 10^, etc. As such neural manifolds capture the potential “states” that the collective activity of a neural population can take.

The investigation of the neural modes and their time-varying activation, or *latent dynamics*, has shed light on questions about the generation of behaviour that had remained elusive when studying the independent function of single neurons. These insights range from behavioural flexibility^11, 12^ and stability^13, 14^, to motor learning^5, 15–19^, principles of overt^20–23^ and covert behaviour^24–27^, and representations of time^28–30^ and space^7, 31^. All these findings are based on the idea that neural manifolds and their latent dynamics capture the functional processes underlying motor control^13, 14, 20, 21^, sensory processing^32–34^—including how these change along the neuraxis^35^—and abstract cognition^7, 8, 31^. Neural manifolds seem to also shape how the brain may deploy new activity patterns to adapt to novel situations^5, 15, 18, 36–38^. Thus, accurately capturing the geometrical properties of neural manifolds is necessary to advance in our understanding of the neural processes underlying behaviour.

Virtually all previous studies adopted linear dimensionality reduction methods such as Principal Component Analysis (PCA) or Factor Analysis to identify the neural modes and their associated latent dynamics from the firing rates of the recorded neural population^39, 40^. As such, it was implicitly assumed that the neural modes capturing the population activity define a lower dimensional surface or *neural manifold*^1, 41^ that is effectively flat—although there are exceptions outside the motor system^7, 8, 31, 42, 43^, the focus of this work. However, the activity of single neurons is inherently nonlinear. For example, at any given moment, a single neuron fires either zero or a positive and finite number of action potentials. Moreover, each neuron makes up to thousands of connections with other neurons, creating intricate connectivity patterns^44–46^. Such intricate connectivity patterns, in turn, should make the interactions between these neurons equally complex, and likely nonlinear. This combination of nonlinear neurons with complex interactions suggests that the neural manifolds underlying the neural population activity during behaviour may be similarly nonlinear (Figure 1A).

**Figure 1:**
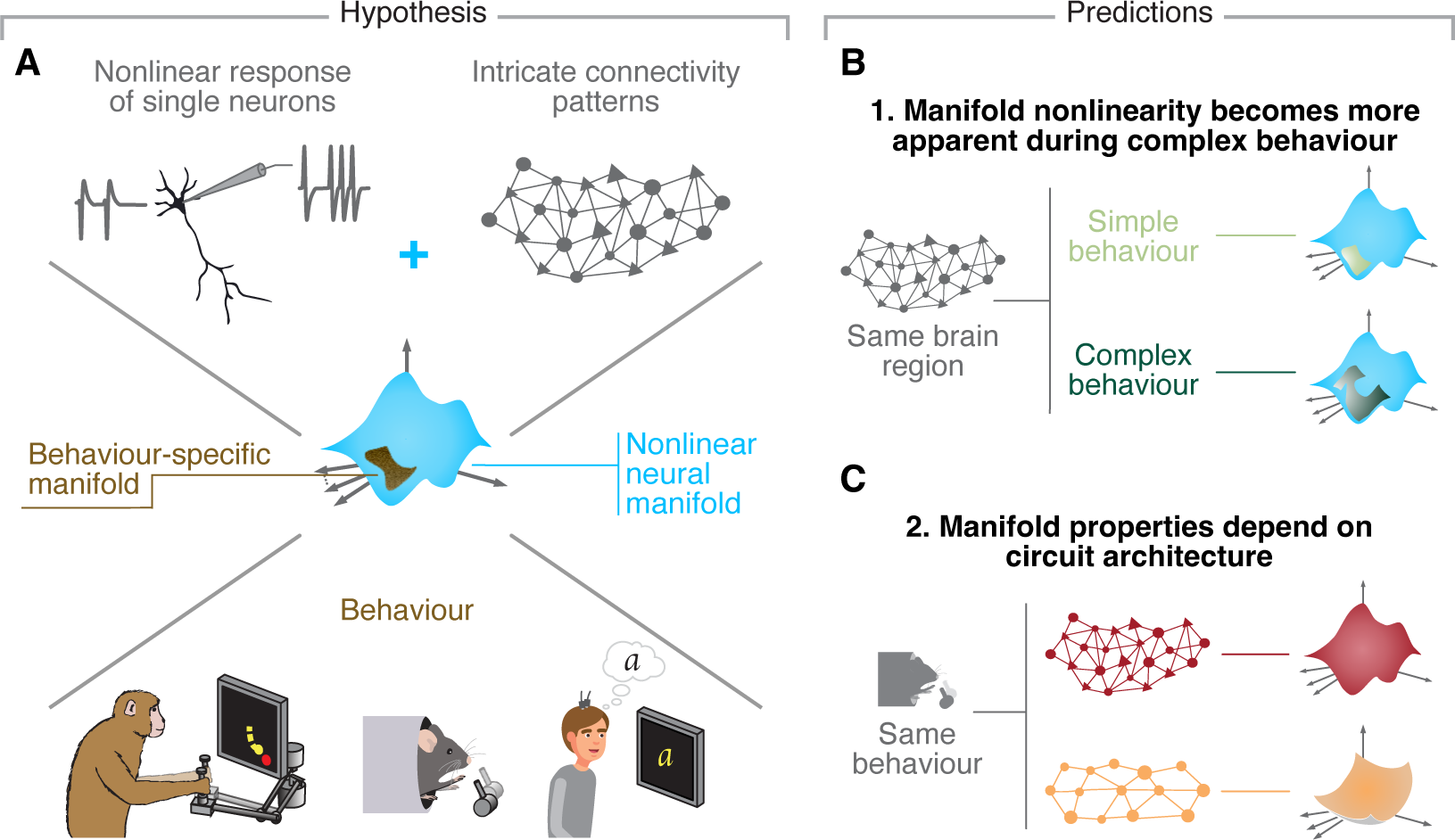
Hypothesis. **A**. Due to the nonlinear activity profiles of single neurons and the complex connectivity profiles of neural circuits, we hypothesized that neural manifolds underlying behaviour shou be nonlinear. **B**. We predicted that more complex behavioural tasks that require a broader range of activity patterns will make the intrinsic nonlinearity of neural manifolds more apparent. **C**. We further predicted that, if the geometry of neural manifolds indeed reflects circuit properties, neural manifolds from cytoarchitecturally different brain regions would exhibit distinct degrees of nonlinearity.

The assumptions underpinning our hypothesis that neural manifolds are intrinsically nonlinear lead to two testable predictions. First, any nonlinear object can be locally approximated by a (hyper)plane. However, this approximation becomes progressively worse—and the nonlinearity more apparent—as more of the surface of this nonlinear object is sampled. Accordingly, looking over a larger extent of the manifold by studying a more complex behaviour will be more revealing of differences between a flat approximation and a more accurate nonlinear description. Thus, we predict that for any given brain region, “complex” tasks that require more varied behaviour and thus elicit a broader range of neural activity patterns should have more apparently nonlinear manifolds than simple tasks that only require a few different activity patterns (Figure 1B). In practice, while in the case of relatively simple, constrained tasks, linear methods may yield reconstructions of the underlying neural manifold that are almost as accurate as those provided by nonlinear methods, we expect them to degrade as task complexity increases. Second, following on our hypothesis that complex circuit connectivity should make neural manifolds nonlinear, we further predict that brain regions with very different circuit architectures should display different degrees of nonlinearity during the same behaviour (Figure 1C). This implies that, in practice, the accuracy of the neural manifold estimates provided by linear methods may be largely different across regions.

We addressed these three hypotheses using a combination of computational models and neural population recordings from monkeys, mice, and humans performing reaching, grasping and pulling, and imagined writing tasks, respectively. We first show that even during a relatively simple centre-out reaching task, the activity of neural populations from monkey motor cortex is better described by a nonlinear rather than a flat manifold. In good agreement with our second hypothesis, the nonlinearity of this neural manifold was much more apparent when considering all eight reach directions as opposed to a single one, a result that we extended by examining population recordings from human motor cortex^22^. There, manifolds underlying a broad range of attempted movements were much more nonlinear than those underlying a more limited repertoire. Next, we addressed our third hypothesis that manifold nonlinearity is determined by circuit properties through comparison of manifolds underlying the activity of simultaneously recorded populations from two cytoarchitecturally distinct motor regions of the mouse brain—motor cortex and the dorsolateral striatum— during a grasping and pulling task^14, 47, 48^. Manifold nonlinearity was indeed markedly different between these two regions, with striatal manifolds being much more nonlinear than motor cortical manifolds. Finally, we used computational models to observe how manipulation of circuit connectivity influences manifold nonlinearity. The nonlinearity of manifolds underlying the activity of recurrent neural network (RNN) models^18, 37, 49–51^ trained to perform the monkey centre-out reaching task was tightly linked to their degree of recurrent connectivity, an observation that held across different network various architectures and hyperparameter configurations. Combined, these results show that intrinsically nonlinear manifolds underlie neural population activity during behaviour, and that the degree of nonlinearity is shaped by both the circuit connectivity and behavioural “complexity”. Considering these nonlinearities will likely be crucial as the field moves toward the study of a broader range of brain regions during ever more complex behaviours.

## Results

### Nonlinear manifolds underlie motor cortical population activity during reaching

We trained two macaque monkeys (C and M) to perform an instructed delay centre-out reaching task using a planar manipulandum (Figure 2A) (Methods). Monkey C performed the task in two sets of experiments, first using the right arm, and then the left arm, which we denote as Monkey C_L_ and C_R_, respectively (L and R refer to the contralateral side of the brain). We recorded neural activity using chronically-implanted microelectrode arrays inserted into the arm area of the primary motor cortex (Monkey C_R_, C_L_, and M) and dorsal premotor cortex (Monkeys C_L_ and M). These recordings, which we combined across areas due to similarities in their activity^52^, allowed us to identify the activity of hundreds of putative single motor cortical neurons during each session (46–290 depending on the session; average, 154±86) (Figure 2B). To better account for the variability in neural activity and behaviour across different trials, we performed all analyses on single trial data rather than on trial averaged data.

**Figure 2:**
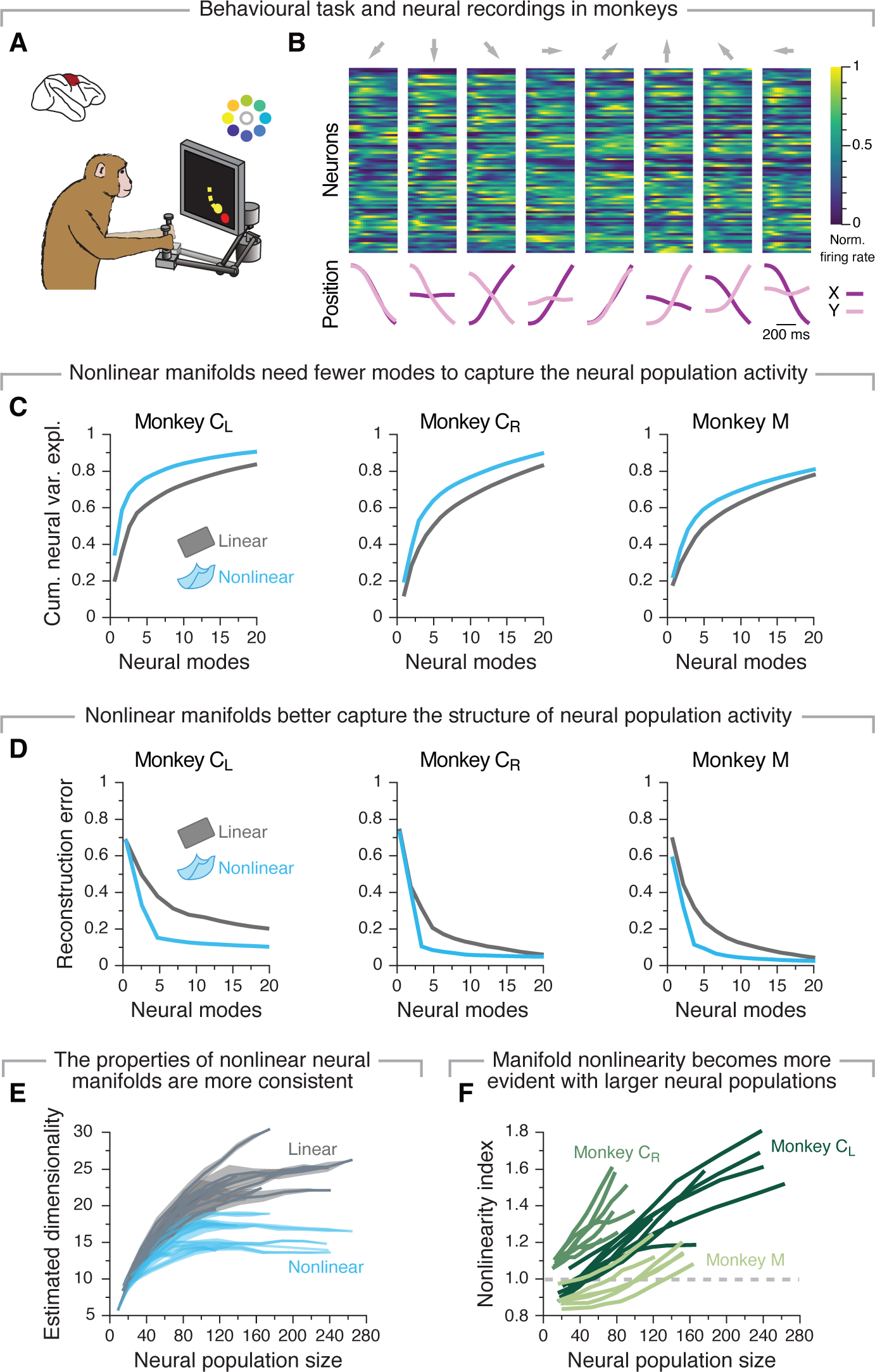
A nonlinear manifold underlies motor cortical population activity during a centre-out reaching task. **A**. Monkeys performed a centre-out reaching task with eight targets. **B**. Average single neuron firing rates and hand positions as a monkey reached to each target. Data from a representative session from Monkey C_R_. **C**. Cumulative neural variance explained by flat (gray) and nonlinear (blue) manifolds as function of the number dimensions. Shown are one example data set for each monkey. **D**. Reconstruction error after fitting flat (gray) and nonlinear (blue) manifolds with increasing dimensionality. Data from the same three example sessions as in C. **E**. Estimated dimensionality of flat (grey) and nonlinear (blue) manifolds as function of the number of neurons used to sample them. Lines and shaded areas, mean s.d. across 10 random subsets of neurons. Data from all sessions from Monkey C_L_. **F**. Nonlinearity index indicating the ratio of the estimated dimensionality of nonlinear manifolds to that of flat manifolds as function of the number of neurons used to sample them. Shown are all sessions from each monkey, colour coded by animal (legend). Values greater than one (dashed grey line) indicate that nonlinear manifolds capture the data better than flat manifolds.

We first identified flat manifolds spanning the neural population activity during movement using PCA^11, 13, 16, 20, 39^. As expected^11, 14, 16, 21, 53^, a large portion of the variance in the neural data was captured by relatively few neural modes (principal components in the case of PCA; gray trace in Figure 2C, and Figure S1). To determine if nonlinear manifolds capture the neural population activity better than flat manifolds, we used a standard nonlinear dimensionality reduction method, Isomap^54^, to find a nonlinear manifold underlying the same neural population activity. As shown in Figure 2C, a nonlinear manifold explained the same amount of variance as a flat manifold with considerably fewer neural modes (compare the blue and gray traces; Figure S1 shows additional examples, Figure S2 validates this approach on known manifolds, and on synthetic neural data^55^). Since explaining more variance with fewer modes is a hallmark of a better representation of the (neural) data, this result indicates that motor cortical manifolds may be intrinsically nonlinear even during a relatively simple task.

We performed additional analyses to verify this observation. First, we assessed whether flat or nonlinear manifolds better capture the structure of neural population activity by quantifying how well their respective latent dynamics can be used to reconstruct the full dimensional activity using a “reconstruction error” metric (called “residual variance” in the original Isomap paper^54^). For all 24 datasets, linear manifolds had considerably larger reconstruction errors than their nonlinear counterparts until at least 10–20 neural modes were considered (Figure 2D; Figure S1; Figure S3A shows that this metric is robust across both different sets of trials and neurons). Moreover, the Isomap reconstruction errors had a clear “elbow”—a dimensionality value at which the error abruptly saturated— indicating that additional nonlinear neural modes did not greatly improve the low-dimensional representation of the neural population activity.

The previous analyses indicate that nonlinear manifolds require fewer dimensions than flat manifolds to capture the variance in neural population activity (Figure 2C), while also allowing for better reconstruction (Figure 2D). As a final analysis to establish the nonlinearity of neural manifolds, we investigated whether the properties of the estimated nonlinear manifold were also more robust against changes in the neurons used for its estimation than those of their linear counterparts. We focused on the estimated dimensionality of the neural manifold and hypothesised that if a method provides a good estimate, using a sufficiently large number of neurons for its estimation should consistently give the same estimated dimensionality (see examples for known manifolds and synthetic data in Figure S2; Figure S3A-C show that this metric is robust against dropping trials and data points). In contrast, if the low-dimensional projection onto the neural manifold fails at fully capturing the structure of the population activity, then the estimated dimensionality will likely increase as more neurons are considered. As expected, for virtually all datasets, the estimated dimensionality of the nonlinear manifolds plateaued after 30–40 neurons were considered (blue traces in Figure 2E; Figure S3D;), whereas that of flat manifolds never reached a plateau even when considering all recorded neurons (65–250). This trend became most apparent when we computed a “nonlinearity index” as the ratio between the estimated dimensionality of the linear and nonlinear manifolds for the same number of neurons. The nonlinearity index mostly took values greater than one and increased monotonically with the number of neurons, indicating that nonlinear manifolds provided progressively better approximations of the neural population activity as the number of neurons increased (Figure 2F). These results were obtained by estimating the manifold dimensionality using the “participation ratio”, defined as the number of neural modes required to explain ~80 % of the total variance^2, 56^, but we obtained similar results when using a different dimensionality estimation metric^55, 57^ (Figure S3E).

We performed additional controls to establish the nonlinearity of neural manifolds. First, we verified that the dimensionality estimates for the linear and nonlinear manifolds were independent of the dimensionality reduction technique used for manifold estimation. Reassuringly, we obtained qualitatively similar results when we used Factor Analysis^5, 58^ instead of PCA to identify the flat neural manifolds (Figure S3F), and when we used nonlinear PCA^59^ instead of Isomap to identify the nonlinear neural manifolds (Figure S3G). Nonlinear manifolds were also more informative about behaviour: decoders trained on the latent dynamics within nonlinear manifolds outperformed those trained on linear manifolds given the same number of neural modes (Figure S3H), indicating that the nonlinearities capture behaviourally relevant information. Taken together, these results suggest that the presence of nonlinear manifolds reflects fundamental features of neural population activity, and not just the greater ability of nonlinear dimensionality reduction methods to fit data. Motor cortical manifolds thus exhibit nonlinear features even for a eight-target centre-out reaching task, and linear approximations of these intrinsically nonlinear structures become progressively more inaccurate as more neurons are considered.

### More varied behaviours reveal greater manifold nonlinearities

We have shown that neural manifolds in monkey motor cortex during a relatively simple centre-out reaching task are nonlinear (Figure 2). Our second hypothesis stated that, during tasks that require a broader range of movements, neural activity would explore a larger portion of the underlying neural manifold thus revealing more clearly its intrinsic nonlinearity (Figure 1B).

To study the relationship between manifold nonlinearity and task complexity, we analysed the activity of neural populations from the “hand knob” area of motor cortex while a paralysed participant attempted to perform a broad range of movements. These included writing straight lines, single letters, and symbols according to a visual cue using their contralateral hand (data from Ref. 22, including ~200 multi-units per session) (Figure 3A,B). We first studied attempted drawing of one-stroke lines of three different lengths across 16 directions. We compared neural manifolds underlying population activity while the participant drew lines in a single direction with those underlying the drawing of lines in all 48 length × direction combinations. First, we observed that the neural manifold during the task of drawing lines in a single direction was rather flat, with both PCA and Isomap giving similar reconstruction error (Figure 3C; Figure S4B), and both of their estimated dimensionalities plateauing before all the neurons in the population were considered (Figure S4C). In contrast, when many line lengths and directions were considered, all our measures indicated that the motor cortical manifold was nonlinear (Figure 3C,D; Figure S4B,C; S4D shows how nonlinearity tends to increase for an increasing number of conditions).

**Figure 3:**
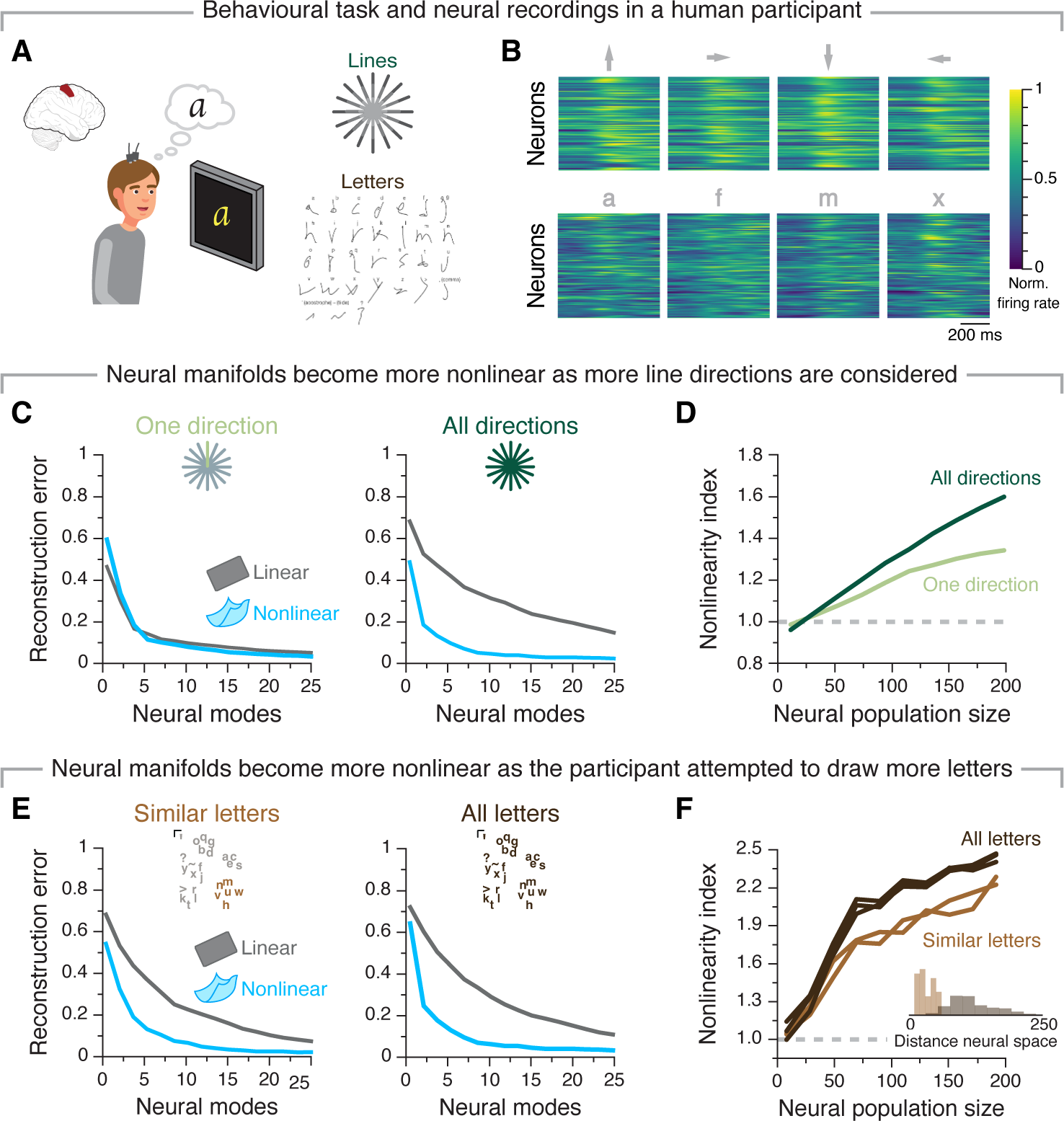
More complex tasks that require more varied neural population activity patterns reveal greater manifold nonlinearity in human motor cortex. **A**. One participant implanted with microelectrode arrays in the hand “knob” area of motor cortex performed a variety of attempted drawing (“Lines task”) and handwriting tasks. **B**. Average firing rates for twelve different letters when attempting to draw lines in four different directions (top) and when attempting to write four different letters (bottom). **C**. Reconstruction error after fitting flat (grey) and nonlinear (blue) manifolds with increasing dimensionality to the neural population activity as the participant attempted to draw strokes in one direction (left) or across all sixteen directions (right). **D**. Nonlinearity index indicating the ratio of the estimated dimensionality of nonlinear manifolds to that of flat manifolds as a function of the number of neurons used to sample them for the simple and complex version of the stroke drawing task. **E**. Same as C but comparing drawing similar letters (left) to drawing all the letters in the English alphabet (right). **F**. Same as D but for the comparison between simple and complex writing tasks. Inset: Neural states associated with attempting to draw letters with similar shapes are closer than those for different letters, as confirmed by an Euclidean distance analysis.

To further investigate this relationship, we re-visited the monkey centre-out reaching task and compared the nonlinearity of neural manifolds underlying reaches to all eight targets to those underlying reaches to one single target (Figure S4G-H). As predicted, the difference in quality of fit between nonlinear and flat neural manifolds was much smaller when only considering a single reaching direction, indicating that manifold nonlinearity increases with task complexity.

We further compared the nonlinearity of neural manifolds underlying even more complex tasks by focusing on the attempted handwriting task. We isolated trials where the participant attempted to draw letters from the English alphabet with very similar shapes^60^, and compared the estimated neural manifolds with those underlying the drawing of all letters. The nonlinear manifolds underlying both tasks were considerably lower dimensional than their flat counterparts (Figure 3E; Figure S4J,K). Moreover, as expected, manifold nonlinearity increased for the larger group of dissimilar letters (Figure 3F), a difference likely driven by the smaller exploration of neural space for the limited set of letters (inset in Figure 3F). The observation that task complexity increases manifold nonlinearity across three different tasks in two different species supports the hypothesis that more complex behaviours that require more varied behaviour make the intrinsic nonlinearity of neural manifolds more apparent.

### Neural manifold nonlinearity changes across cytoarchitecturally distinct brain regions

Our third hypothesis that manifold nonlinearity is shaped by network connectivity properties implies that different brain regions may be better or worse approximated by flat manifolds depending upon their circuit properties. To address this presumed relationship *in vivo*, we compared the nonlinearity of motor cortical manifolds to another area critical for forelimb movement but with very different microarchitecture and cell types: the dorsolateral striatum (henceforth just striatum). For example, while motor cortex has more recurrent excitatory connectivity, and has a broad range of cellular nonlinearities^61, 62^ (due to large dendritic tree and many interneuron types), striatum, does not present these different neuron types, and has a very different circuitry dominated by feedforward and recurrent inhibitory connectivity^63, 64^.

We analysed simultaneous motor cortical and striatal population recordings from four mice performing a reaching, grasping and pulling task^14, 47^(Methods; Figure 4A,B) (motor cortical neurons: 54–96 depending on the session; average, 76 15, mean s.d.; striatal neurons: (62–106 depending on the session; average, 83 14, mean s.d.). A direct comparison of the nonlinearity of motor cortical and striatal manifolds provided direct evidence that manifold nonlinearity can be strikingly different across different brain regions: all of our three measures revealed that striatal manifolds are nonlinear whereas motor cortical manifolds estimated during this behaviour appear mostly flat (Figure 4C,D; Figure S5A,B, S6). This was the case even if single neuron firing rate statistics were similar between the two regions (Figure S5C), and when considering that the nonlinear aspects of striatal neuron biophysics are less marked than those of motor cortical neurons^62, 65^. Thus, our experimental findings suggest that circuit properties may indeed be the primary factor underlying neural manifold nonlinearity.

**Figure 4:**
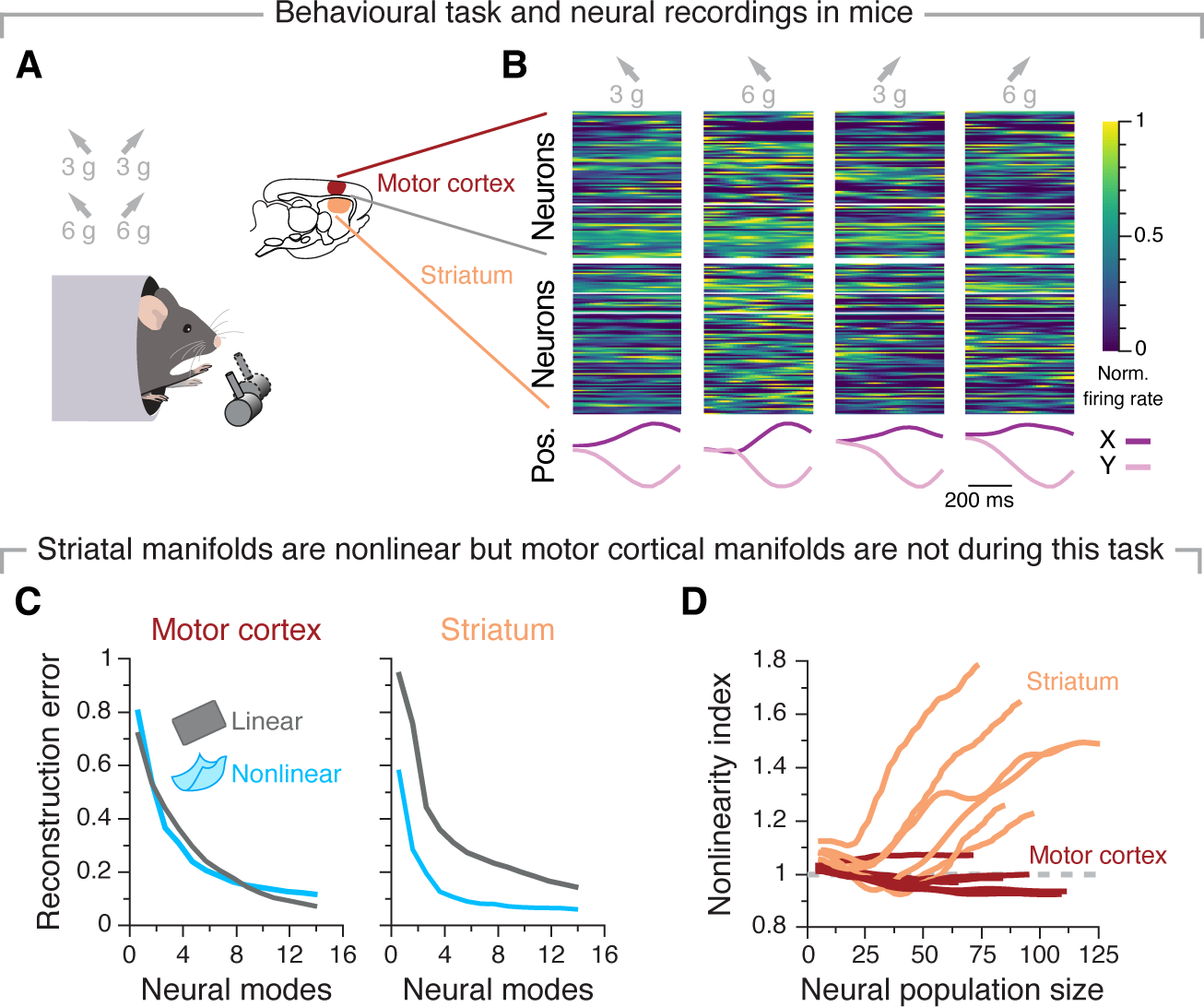
The nonlinearity of neural manifolds changes across architecturally distinct brain regions. **A**. Mice performed a reaching, grasping, and pulling task with four conditions (two positions two targets). **B**. Average single neuron firing rates for motor cortex (top) and striatum (bottom), and hand positions as one mouse performed each condition. **C**. Reconstruction error after fitting flat (grey) and nonlinear (blue) manifolds with increasing dimensionality to the motor cortical (left) and striatal (right) population activity. Data from the same example mouse as in B. **D**. Nonlinearity index indicating the ratio of the estimated dimensionality of nonlinear manifolds to that of flat manifolds as function of the number of neurons used to sample them. Shown are all mouse sessions, colour coded by region (legend). Values greater than one (dashed grey line) indicate that nonlinear manifolds capture the data better than flat manifolds.

### A neural network model to understand the emergence of manifold nonlinearity

The previous analyses provided support for our hypotheses that neural manifolds are intrinsically nonlinear (Figure 2,3), and manifold nonlinearity is influenced by circuit connectivity properties (Figure 4). Since synaptic connectivity could not be manipulated during our behavioural experiments, here we investigated directly the relationship between manifold nonlinearity and circuit connectivity by developing RNN models of motor cortical activity with different structure in their recurrent weights. We trained RNNs to perform the same instructed delay reaching task in monkeys (Methods) that was used to demonstrate our core experimental results. Our model architecture was based on previous studies using RNNs to simulate motor cortical activity^18, 37, 49–51^, but adapted to account for trial-to-trial variability during reaches to the same target. We achieved this by training RNNs to produce the monkey’s hand velocity during each recorded trial using the actual trial-specific preparatory neural activity as input, rather than the typical target cue (Methods; Figure 5A,B; Figure S7A,B). RNNs trained with this approach recapitulated key features of the actual neural activity, both at the population level (Figure 5C; Figure S7D) and the single unit level, including clear—albeit more moderate—fluctuations across different trials to the same target (Figure S7C). These observations established our RNNs as suitable models for the actual neural activity.

**Figure 5:**
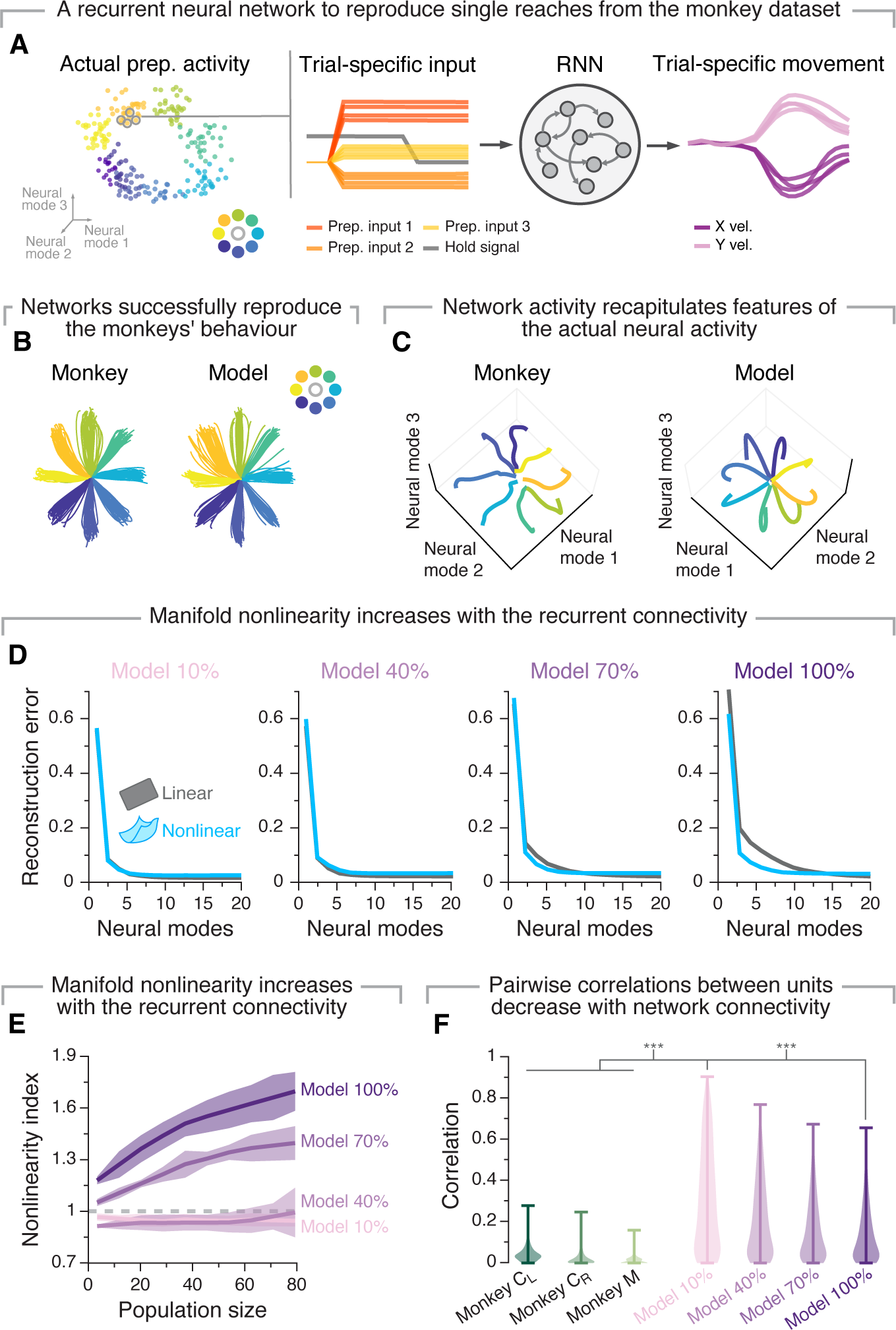
A recurrent neural network model supports the relationship between circuit connectivity and manifold nonlinearity. **A**. We trained recurrent neural network models to generate the actual hand velocities generated by the monkeys using trial-specific preparatory activity as inputs. **B**. These models reproduced the variability across reaches to the same target observed experimentally. **C**. The latent dynamics of the network activity recapitulated the structure of the monkey latent dynamics (systematic quantifications in Figure S7). Individual traces, trial averaged activity for each target. **D**. Reconstruction error after fitting flat (grey) and nonlinear (blue) manifolds with increasing dimensionality. Data for example networks with increasing degrees of recurrent connectivity (10%, 40%, 70% and 100%). **E**. Nonlinearity index indicating the ratio of the estimated dimensionality of nonlinear manifolds to that of flat manifolds as function of the number of sampled units. We compare results for networks with increasing degrees of recurrent connectivity (legend). Values greater than one (dashed grey line) indicate that nonlinear manifolds capture the data better than flat manifolds. Lines and shaded areas, mean s.d. across 15 repetitions for each of the 10 seeds. **F**. Strength of pairwise correlations between units from networks with different degrees of recurrent connectivity, compared to the actual monkey data. Violin, probability density for each network connectivity and monkey level. \*\*\**P <* 0.001, two-sided Wilcoxon rank-sum test.

We next investigated the relationship between the degree of recurrent connectivity of the network and manifold nonlinearity. We originally hypothesized that manifold nonlinearity is in part due to the extensive number of connections across brain neurons. Consequently, we predicted that for networks with the same architecture manifold nonlinearity would increase with the degree of recurrent connectivity. Our results show that the population activity for networks with low connectivity (10% and 40%) was spanned by a relatively flat manifold, as indicated by the similarity in variance explained (Figure S7E), reconstruction error (Figure 5D) and estimated dimensionality (Figure S7F) between flat and nonlinear manifolds. High connectivity networks (70% and 100%), in contrast, had manifolds that were clearly nonlinear based on all three metrics (Figure 5D; Figure S7E,F). Most notably, the dimensionality of nonlinear manifolds reached a plateau after relatively few units, while that of the flat manifolds continued to increase as more units were considered (Figure S7F). This resulted in nonlinearity indexes presenting values well above one (Figure 5E). These differences were driven by differences in recurrent connection probability not the strength of the recurrent weights, since the weight distributions were similar across all trained networks (Figure S7G). Therefore, when considering these “standard” RNN models, only the activity of networks with dense recurrent connections was spanned by nonlinear manifolds, with manifold nonlinearity increasing with the level of recurrent connectivity. This trend held for a different class of recurrent network models (Figure S9), and even when we trained RNNs with linear rather than nonlinear units to perform this same task (Figure S8A). In stark contrast, fully connected feedforward networks that lacked recurrent connectivity had flat manifolds (Figure S8B). Combined, these results suggest that dense recurrent connectivity may be both necessary and sufficient for neural manifolds to become nonlinear.

While the RNN models above exhibited clear similarities to the recorded neural activity (Figure 5; Figure S7), the pairwise correlations between units were much larger than those observed experimentally^34, 66–68^ (compare “Monkey” and “Model” distributions in Figure 5F). Yet, intriguingly, increasing the degree of recurrent connectivity, which brought manifold nonlinearity closer to the larger experimentally observed values (Figure 2), also decreased moderately the pairwise correlations between units. This inverse relationship between manifold nonlinearity and pairwise correlations between units also present in the mouse recordings—striatum showed lower pairwise correlation between units (Figure S5D) and higher manifold nonlinearity than motor cortex (Figure 4C-D)—suggesting a potential fundamental association between these two measures.

We explored this relationship by training a new set of RNNs with the additional constraint of producing lower pairwise unit correlations (Methods; Figure 6A) to test more directly the association between these metrics. As expected, while these “decorrelated” networks also learned the task successfully (Figure 6B; Figure S10A) and produced activity patterns similar to those observed in monkey motor cortex (Figure S10D), the resulting unit correlations were much closer to the experimentally observed correlations than those of our previous standard models (Figure 6C; Figure S10B). The manifolds of the decorrelated networks were much more nonlinear than those of the standard networks, even for matching degrees of recurrent connectivity (Figure 6D,E; Figure S10E), while still exhibiting an association between recurrent connectivity level and manifold nonlinearity (Figure S10C). In agreement with our hypothesis that circuit connectivity shapes manifold nonlinearity, the differences in pairwise correlations and manifold nonlinearity between standard and decorrelated networks corresponded to differences in their connectivity: overall, the weight changes required by the standard networks to learn the task were higher dimensional than those of decorrelated networks (Figure S10G), even if their distributions did not exhibit striking differences (compare Figure S7G and S10F). Combined, our simulations provide strong support for our experimental observation that network connectivity is indeed an important factor shaping manifold nonlinearity. While the activity of sparsely connected standard networks could be captured by a flat manifold, the more physiologically relevant condition of dense recurrent connectivity led to the emergence of nonlinear manifolds.

**Figure 6:**
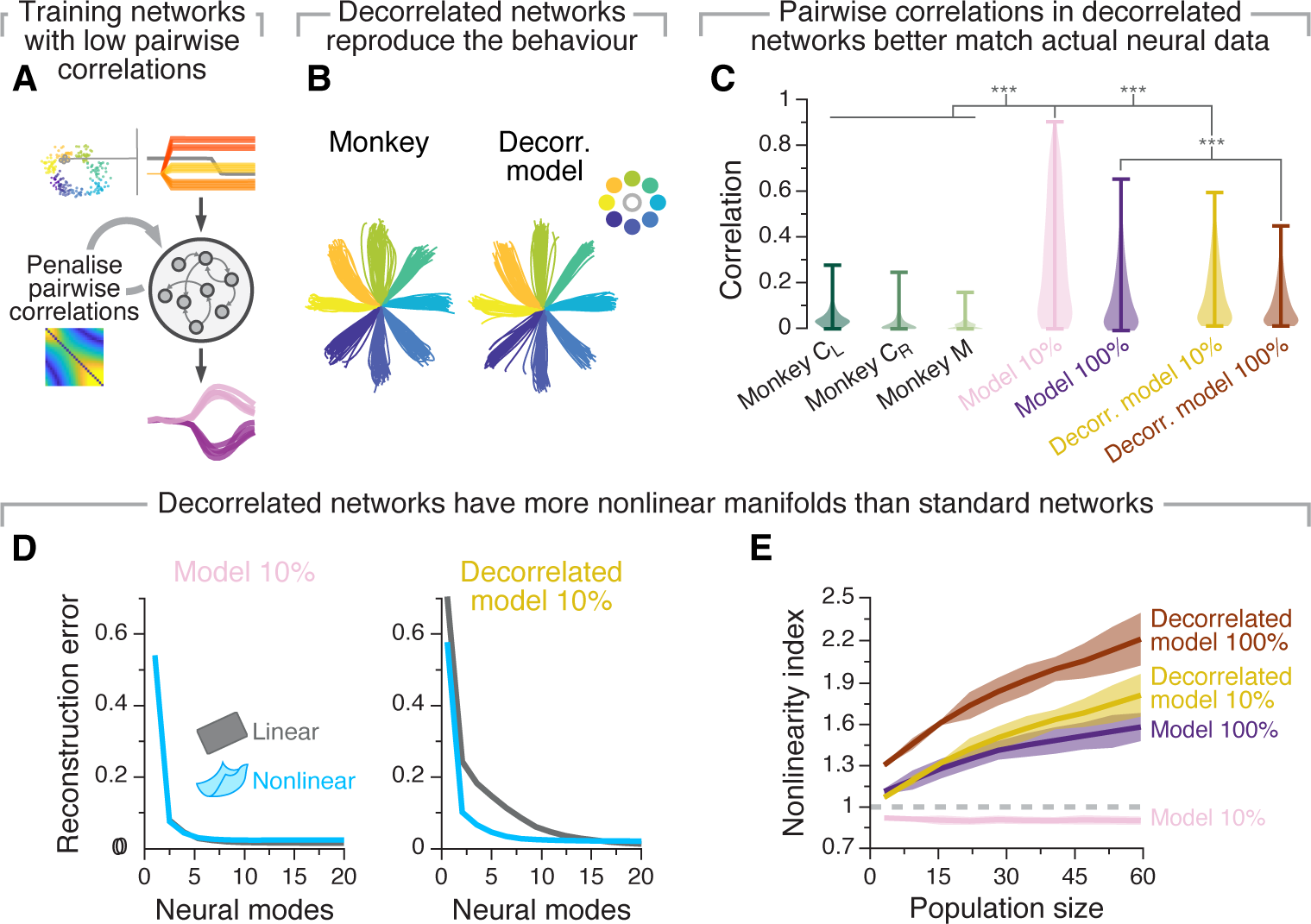
A recurrent neural network model constrained to have low pairwise correlation between units has more nonlinear manifolds. **A**. We modified the training procedure of our models so as to limit their pairwise correlations between units to better match experimental neural data. **B**. These models also reproduced the experimentally recorded hand trajectories. **C**. “Decorrelated networks” have pairwise correlations between units that are similar in magnitude to those experimentally observed in monkeys. Shown are the strength of pairwise correlations between units from “decorrelated networks” with two degrees of recurrent connectivity compared to “standard networks” with similar degrees of connectivity, along with the pairwise correlations experimentally observed in monkeys. Violin, probability density for each network connectivity level and monkey. \*\*\**P <* 0.001, two-sided Wilcoxon rank-sum test. **D**. Reconstruction error after fitting flat (grey) and nonlinear (blue) manifolds to “standard” (left) and “decorrelated” (models) with similar levels of recurrent connectivity (10%). **E**. Nonlinearity index indicating the ratio of the estimated dimensionality of nonlinear manifolds to that of flat manifolds as function of the number of sampled units. We compare results between various standard and decorrelated networks (results for additional degrees of recurrent connectivity are shown in Figure S10C). Lines and shaded areas, mean*±*s.d. across seeds.

## Discussion

Technological, computational, and theoretical advances have fostered an expansion from the study of single-neuron activity and the encompassing neural circuits to the investigation of the latent dynamics reflecting the coordinated activity of neural populations^1, 41, 69, 70^. In practice, these latent dynamics are often inferred by projecting the recorded neural activity onto a neural manifold that is identified using a particular dimensionality reduction or manifold learning method^39, 40^. As such, the particular choice of methodology translates into implicit assumptions about the properties of the neural manifold. Most studies have relied on linear methods such as PCA or Factor Analysis to identify the latent dynamics, thereby assuming that the neural population activity lies in a flat neural manifold. Here, we tested the hypothesis that neural manifolds underlying behaviour are intrinsically nonlinear by comparing the relative performance of linear and nonlinear dimensionality reduction methods on recordings from a variety of tasks, brain regions, and species. We showed that, in virtually all cases, the activity of neural populations from motor regions of the brain is best described by a nonlinear rather than a flat manifold. This difference between linear and nonlinear methods became more dramatic as task complexity (e.g., number of different movements) increased. During the same task, architecturally distinct brain regions differed in the degree of nonlinearity of their manifolds, an observation that we further supported by manipulating network connectivity in RNNs performing the same task.

### Influence of task complexity on manifold nonlinearity

The nonlinearities inherent in neural manifolds became more apparent as the complexity of behaviours increased, an observation that held for the motor cortex in both human (Figure 3) and monkey (Figure S4E,F). Intuitively, we proposed that this can be explained by simple behaviours exploring only a small portion of the available neural states. Even if the “landscape” defined by these states were intrinsically nonlinear, these small explorations could be well approximated locally by a flat manifold. However, as the complexity of behaviours increases, activity explores a larger region of state space and in the process makes manifold nonlinearities more apparent (Figure 1B).

A previous theoretical study on the properties of flat motor cortical manifolds proposed that the upper bound on their dimensionality should increase with task complexity^2^. Our results suggest an alternative explanation: increasing the number of movements an animal performs may make the neural activity explore a larger portion of an intrinsically nonlinear manifold. This should in turn increase the difference in the number of dimensions needed by linear and nonlinear manifolds to accurately capture the neural population activity^3, 71, 72^.

Our data from monkey and mouse motor cortex seemed to indicate an apparent difference between these two species: during the studied motor tasks, monkey motor cortical manifolds were nonlinear (Figure 2) while mouse motor cortical manifolds were flat (Figure 4). The differences in cytoarchitecture between these two species^73^ could be one factor driving this difference, but our data suggest that behavioural complexity is the primary cause. The monkeys reached to eight different targets by producing a relatively large variety of muscle activation patterns^74, 75^, including arm flexion and extension. The mice, in contrast, reached in only one direction–away from the body^14, 47^. Thus, the mice performed a far simpler behaviour and thus their neural activity may have explored only a small portion of the motor cortical state space. This interpretation is supported by our direct comparison between reaches to all eight different targets and reaches to a single target during the monkey centre-out reaching task (Figure S4D-F). In the latter case, which is more akin to the mouse dataset, manifolds were almost flat, as the nonlinearity indices became similar (Figure S4D).

For practical and conceptual reasons, most neuroscientists study relatively low-dimensional behaviours produced by animals with extensive practice in a given task. Linear methods that estimate a flat manifold defined based on a series of orthogonal directions may be the best approach for this kind of tasks, especially given that geometric interpretation may be more intuitive. However, our results indicate that as the field moves toward the study of more complex naturalistic behaviours (e.g., Ref. 76–78), we will need to consider nonlinear aspects of manifold geometry. This poses the additional challenge of choosing one among the many nonlinear dimensionality reduction or manifold learning methods. In practice, since each method has its inherent assumptions and these may lead to different geometries (and distortions), e.g., based on how well they preserve global vs. local features^40^, one may need to try several methods to achieve an accurate view of a manifold’s geometry, especially when *a priori* predictions are not available. Defining the dimensionality of behaviour also remains a looming challenge. Here, we have assumed that tasks requiring more varied outputs—be it intended, as in the case of the human BCI participant (Figure 3), or actual, as in the case of the monkeys (Figure S4E,F)—are more complex. While this assumption is reasonable for the present study, a recent proposal to define the dimensionality of behaviour based on the number of past features that maximally predict future movements^79^ could establish a more rigorous relationship between the dimensionalities of the neural manifold and behavior.

### Factors driving the region-specificity of manifold nonlinearity

We investigated possible biophysical factors leading to the intrinsic nonlinearity of neural manifolds. We predicted that circuit connectivity would be a key determinant of manifold nonlinearity, and found that increasing recurrent connectivity monotonically increased manifold nonlinearity for two different artificial neural network architectures (Figure 5; Figure S7, S9). Moreover, flat manifolds accurately captured the latent dynamics for only a subset of sparsely connected networks (Figure 5; Figure S9). This association between recurrent connectivity and manifold nonlinearity held for RNN models with both nonlinear and linear units (Figure S8A) but was absent in purely feedforward fullyconnected networks with nonlinear units (Figure S8B). Thus, manifold nonlinearity may result not from the properties of the individual units, but rather from the interactions among the constituent units in an emergent fashion^80, 81^.

We further confirmed the link between circuit connectivity and manifold nonlinearity experimentally. Neural manifolds from the architecturally distinct mouse motor cortex and dorsolateral striatum showed different degrees of nonlinearity during the same behaviour even while providing comparable predictions of movement kinematics^14, 47^. Based on our RNN results, we argue that this difference arises at least in part from their distinct cytoarchitecture^61–64^. Yet, the difference in manifold nonlinearity between motor cortex and striatum could also be driven by differences in the types of inputs they receive. Indeed, compared to M1, striatum may integrate inputs from a wider variety of brain regions, encompassing motor, associative, sensory, and limbic regions^63, 82, 83^. Integration of such varied information streams may lead to more intricate population-wide activity patterns that are better approximated by a nonlinear manifold.

Both our RNN models and the comparison between mouse motor cortex and striatum showed that more nonlinear manifolds are associated with lower pairwise correlations between neurons (Figure 5F and 4D, respectively). Moreover, when we limited the value of these correlations during training, RNNs with comparable degrees of recurrent connectivity exhibited stronger manifold nonlinearities (Figure 6). Interestingly, a previous study linked lower pairwise correlations to a higher dimensionality in flat manifolds^84^. Our results suggest that the increase in dimensionality of the embedding flat manifold may be driven by an increase in nonlinearity, perhaps without a comparably large increase in the intrinsic dimensionality^85^. That is, the number of variables needed to characterise the neural population activity may remain constant even if the pairwise correlations between neurons and the dimensionality of the flat “embedding” manifold increases.

### Manifold nonlinearity beyond the motor system

While we have largely focused on regions in the motor system in the present work, we believe that nonlinear manifolds may be an ubiquitous feature of neural population activity throughout the brain. Yet, the different roles of different systems may translate into fundamental differences in the geometric and topological properties of these underlying nonlinear manifolds. For example, primary sensory regions process rapidly changing stimuli, and are thus likely more input-driven than motor regions controlling smooth limb movements. A recent study analysing the responses of large neural populations in mouse primary visual cortex (V1) found that the dimensionality of flat manifolds increases with the number of visual stimuli^32^, a trend that parallels our findings in human (Figure 3) and monkey (Figure S4E,F) motor cortex. Our results suggest that these V1 manifolds with increasing dimensionality may actually approximate a much lower-dimensional neural manifold that is strikingly nonlinear. Relatedly, a recent study of the mouse whisker system suggests that responses of somatosensory populations are best described by nonlinear manifolds^33^, providing evidence of nonlinear manifolds in primary somatosensory regions. The amount of nonlinearity likely varies as one moves further from primary sensory organs, consistent with the recent observation of a gradient in linear manifold dimensionality along the visual system^86^.

“Higher” brain regions that are less directly linked to the production of behaviour or sensory processing seem to also have neural manifolds with complex and interesting nonlinear geometries. For example, the activity of populations of head-orientation cells in the thalamus lie in a ring-shaped manifold the coordinates of which map onto the animal’s heading direction^8^. Similarly, the activity of enthorinal “grid cells” lies in a toroidal manifold that forms a tessellated, robust, and accurate representation of the environment^7^. Both these manifolds are preserved between awake behaviour and different phases of the sleep cycle^7, 8, 87^, suggesting that their features may be dominated by biophysical constraints (e.g., circuit properties) on neural population activity that are invariant to behavioural states. This conservation of manifold geometry across behavioural states seems at odds with the changes reported for the motor system, where producing different behaviours leads to dramatic changes in the orientation of flat manifolds^12^—although these changes are absent when animals produce various related behaviours^11^.

Finally, the activity of neural populations in the hippocampus, a brain region involved in representing abstract maps of concepts^88^ including space^89^, also lies on a nonlinear manifold whose geometry seems to be flexibly shaped by experience^31, 90^. Indeed, hippocampal manifold geometry changes as animals familiarise themselves with a new environment^90^, or when they link their spatial maps to other cognitive variables, such as value^31^. Therefore, growing evidence suggests that nonlinear neural manifolds may be a universal feature across many cytoarchitecturally distinct regions in the brain.

### Conclusion

Investigating the coordinated activity of neural populations has furthered our understanding of how the brain generates behaviour. However, leveraging this approach to understand more naturalistic behaviours will likely depend upon adequately estimating the neural manifolds underlying the population activity. Here, we have shown that, during a variety of motor tasks, neural manifolds are intrinsically nonlinear, their degree of nonlinearity varies across cytoarchitecturally different brain regions, and becomes more evident during more complex behaviours. These results extend recent reports of nonlinear manifolds across a variety of non-motor regions of the brain^7, 8, 28, 31^, to which we expect our findings to also translate.

From a translational point of view, accounting for the nonlinear geometry of neural manifolds across the sensorimotor system may be key to develop brain-computer interfaces that restore function across a broad range of behaviours based on “decoding” control signals from brain activity^91, 92^. From a fundamental science point of view, accounting for the region-specific nonlinearity of neural manifolds, which is present in cortical and subcortical regions during both overt and covert behaviour, may be crucial to understand the neural basis for more complex and naturalistic behaviours.

## Methods

### Subjects and tasks

#### Monkey

Two monkeys (both male, Macaca mulatta; Monkey C, 6–8 years during these experiments, and Monkey M, 6–7 years) were trained to perform a standard centre-out reaching task using a planar manipulandum. They both had performed a similar task for several months prior to the neural recordings, so they were proficient at it. In this task, the monkey moved their hand to the centre of the workspace to begin each trial. After a variable instructed delay period (0.5–1.5 s), the monkey was presented with one of eight outer targets, equally spaced in a circle and selected randomly with a uniform probability. Then, an auditory go cure signalled the animals to reach the target. The trial was considered successful if the monkey reached the target within 1 s after the go cue, and held the position for 0.5 s. As the monkey performed this task, we recorded the position of the endpoint of the manipulandum at a sampling frequency of 1 kHz using encoders in each joint, and digitally logged the timing of task events, such as the go cue. Portions of these data have been previously published and analysed in Ref. 13, 16, 93 among others, and are publicly available on Dryad (https://doi.org/10.5061/dryad.xd2547dkt).

#### Mouse

After habituation to head-fixation and the recording setup, four 8–16 week old mice were trained to reach, grasp, and pull a manipulandum (similar to the tasks in Ref. 48, 94) for approximately one month. In this task, mice had to reach and pull a joystick positioned approximately 1.5cm away from the initial hand position. The joystick was placed in one of two positions (left or right, ¡ 1 cm apart), and was weighed with two different loads (3 g or 6 g), adding to a total of four trial types. During the experiments, mice could self-initiate a reach to the joystick, followed by the inward pull to get a liquid reward, which was delivered ~1 s after pull onset in successful trials only. A minimum inter-trial period of 7 s was required. Each trial type was repeated 20 times before the task parameters were switched to the next trial type without any cue. For each session, there were two repetitions of each set of four trial types, presented in the same order, making up 2 *×* 4 *×* 20 = 160 trials.

Two high-speed, high resolution monochrome cameras (Point Grey Flea3; 1,3 MP Mono USB3 Vision VITA 1300; Point Grey Research Inc., Richmond, BC, Canada) with 6-mm to 15-mm (f/1.4) lenses (C-Mount; Tokina, Japan) were placed perpendicularly in front and to the right of the animal. A custom-made near-infrared light-emitting diode light source was mounted on each camera. Cameras were synced to each other and captured at 500 frames/s at a resolution of 352 by 260 pixels. Video was recorded using custom-made software developed by the Janelia Research Campus Scientific Computing Department and IO Rodeo (Pasadena, CA). This software controlled and synchronized all facets of the experiment, including auditory cue, turntable rotation, and high-speed cameras. Fiji video editing software was used to time stamp in the videos. Annotation of behavior was accomplished using Janelia Automatic Animal Behavior Annotator^95^ (JAABA).

#### Human

We analysed publicly available data by Willett et al.^22^ (https://doi.org/10.5061/dryad.wh70rxwmv). This data was recorded while a BrainGate study participant (T5) attempted to write a variety of digits, letters, and symbols. Participant T5, a 65 year old man at the time of data collection, with a 4 AIS C (ASIA Impairment Scale C – Motor Incomplete) spinal cord injury that occurred approximately 9 years prior to study enrollment. As a result from this injury, his only hand movements were limited to twitching and micromotions. Among the three tasks in this dataset, we analysed the two that allowed us to most consistently explore a broad range of attempted movements, which we hypothesised would reveal an increase in manifold nonlinearity. These were: 1) attempting to draw straight lines of three different lengths across 16 different directions using a single pen stroke; and 2) attempting to write one out of all the letters and symbols of the English alphabet.

In both types of tasks, Participant T5 was presented with visual cues displayed on a computer monitor. Each trial began with an instructed delay period of variable length (2.0–3.0 s), during which a single character appeared on the screen above a red square that served as hold cue. During the delay period, T5 waited and prepared to attempt drawing the appropriate stroke or writing the corresponding character. Then, the red square in the center of the screen turned green, instructing T5 to begin trying to attempt drawing the stroke or writing the character. After the 1 s go period, the next trial’s instructed delay period began.

### Neural recordings

#### Monkey

All surgical and experimental procedures were approved by the Institutional Animal Care and Use Committee of Northwestern University under protocol #IS00000367. For recording, we used 96-channel Utah microelectrode arrays implanted in the primary motor cortex (M1) and dorsal premotor cortex (PMd) using standard surgical procedures. When recordings of both areas were available, these were pooled together and were denoted as motor cortex. Implants were located in the opposite hemisphere of the hand the animal was using in the task. Monkey C received two sets of implants: a single array in the right M1 while performing the task with the left hand; and, later, two arrays in the left hemisphere (M1 and PMd) while using the right hand for the task. These sessions are denoted C_R_ and C_L_ respectively. Monkey M received dual implants in the right M1 and PMd. We analysed data from seven sessions from each of the three monkeys.

Neural activity was recorded during the behaviour using a Cerebus system (Blackrock Microsystems). The recordings on each channel were band-pass filtered (250 Hz – 5kHz), digitised (30 kHz) and then converted to spike times based on threshold crossings. The threshold was set to 5.5 the root-mean square activity on each channel. We manually spike sorted all the recordings to identify putative neurons (Offline Sorter v3, Plexon, Inc, Dallas, TX). Overall, we identified an average of 278 ± 33 neurons during each session from monkey C_L_ (range, 207–309), 72 ± 17 neurons during each session from monkey C_R_(range, 46–93), and 125 ± 23 neurons during each session from monkey M (range, 92–159).

#### Mouse

All surgical and experimental procedures were approved by the Institutional Animal Care and Use Committee of Janelia Research Campus. A brief (*<*2h) surgery was first performed to implant a 3D-printed headplate^96^. Following recovery, the water consumption of the mice was restricted to 1.2 ml per day, in order to train them in a behavioural task. Following training, a small craniotomy for acute recording was made at 0.5 mm anterior and 1.7 mm lateral relative to bregma in the left hemisphere. A Neuropixels probe was centred above the craniotomy and lowered with a 10 degree angle from the axis perpendicular to the skull surface at a speed of 0.2 mm/min. The tip of the probe was located at 3mm ventral from the pial surface. After a slow and smooth descent, the probe was allowed to sit still at the target depth for a least 5 min before initiation of recording to allow the electrodes to settle. We analysed data from two sessions from each of mouse 38 and mouse 40, and one session from each of mouse 39 and mouse 44.

Neural activity was filtered (high-pass at 300 Hz), amplified (200× gain), multiplexed, and digitised (30 kHz), and recorded using the Spike GLX software (https://github.com/billkarsh/SpikeGLX). Recorded data were pre-processed using the open-source software Kilosort 2.0 (https://github.com/MouseLand/Kilosort), and manually curated using Phy (https://github.com/cortex-lab/phy) to identify putative single units in each of the primary motor cortex and dorsolateral striatum. A total of six experimental sessions (from four different mice) with simultaneous motor cortical and striatal recordings were included in this work. We identified an average of 76 *±* 15 M1 neurons (range, 54–96), and 83 *±* 15 striatal neurons (range, 62–106) in each session.

#### Human

Participant T5 was implanted with two 96-channel Utah microelectrode arrays implanted in the hand “knob” area of the precentral gyrus. The publicly available neural data includes “multi-unit activity”, consisting of binned spike counts (10 ms bins) indicating the number of times the voltage time series on a given electrode crossed a threshold set to 3.5 the root-mean square activity on that channel for 192 channels per session. We analysed data from a total of three sessions.

## Data analysis

We used a similar approach to analyse the monkey, mouse, and human data. First we discarded all the unsuccessful (in the case of the animals, unrewarded) trials. An equal number of trials to each target was randomly selected (eight targets for the monkey and four conditions for mice). We focused our analyses on the following analysis windows: for the monkey data, -100 – 400 ms with respect to the go cue; for the mouse data, -100 – 400 ms with respect to movement onset; and for the human data, -100 – 600 ms with respect to the go cue.

Before performing any dimensionality reduction analysis to identify the underlying linear or nonlinear neural manifold, we first computed the smoothed firing rates as a function of time for each single neuron (or multi-units, in the case of the human data). We obtained these smoothed firing rates by applying a Gaussian kernel (*σ* = 50 ms) to the binned square-root transformed firings (bin size = 20 ms) of each single unit or multi-unit (henceforth, simply “units”). We only excluded units with a low mean firing rate (*<* 1 Hz mean firing rate across all time bins).

### Estimating flat and nonlinear manifolds using linear and nonlinear dimensionality reduction techniques

Following the pre-processing described in the previous section, we concatenated all the trials, producing a neural data matrix **X** of dimension *n* by *T*, where *n* is the number of units and *T* is the total number of time points from all trials. Note that for each session all trials were truncated to the same duration, and thus *T* equals the number of trials × the duration of each trial. For each session, the simultaneous activity of all *n* recorded units was represented in a neural state space. In this space, the joint recorded neural activity at each time bin is represented as a single point, the coordinates of which are determined by the firing rate of the corresponding units. As the activity evolves over time, this point describes a trajectory in neural state space, the *latent dynamics*. We estimated the linear and nonlinear manifolds underlying this neural population activity to understand whether nonlinear manifolds outperform their flat counterparts at capturing its properties. We used Principal Component Analysis (PCA) to compute flat manifolds and Isomap^54^ to compute nonlinear manifolds, although we later replicated the main analysis using different linear and nonlinear dimensionality reduction methods (see section “Additional analyses including controls” below).

PCA is a linear technique for dimensionality reduction that identifies an set of orthogonal directions (eigenvectors) that capture the greatest variance in the data. These directions are ranked based on the amount of variance they explain, which is quantified by the associated eigenvalue. In contrast, Isomap is a nonlinear dimensionality reduction technique that finds a nonlinear manifold that embeds the data. The Isomap algorithm starts by constructing a neighborhood graph based on the pairwise distances between data points. It connects each data point to its nearest neighbors, forming a graph where the distances between connected points are approximately preserved. Next, Isomap estimates the pairwise geodesic distances between all data points on the graph. Once the geodesic distances are computed, Isomap employs classical multidimensional scaling to embed the data points into a lower-dimensional manifold. We refer to the dimensions identified by any dimensionality reduction method simply as *neural modes*.

We used three different measures to establish how well flat and nonlinear manifolds captured the neural population activity. First, we calculated the total neural variance explained by manifolds with increasing dimensionality by computing the cumulative sum of the eigenvalues of the covariance matrix (for PCA) and the double-centered geodesic distance matrix (for Isomap) (see, e.g., Figure 2C). In both cases, the eigenvalue corresponding to a given dimension was divided by the total sum of the eigenvalues, bounding the values between zero and one.

As a second measure, we computed the reconstruction error, RE, provided by each method^54^:

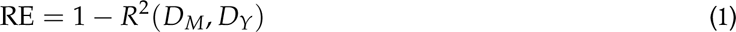

where *D_Y_* is the matrix of Euclidean distances in the low-dimensional embedding found by each method, and *D_M_* is each method’s best distance estimate; in the case of Isomap, this corresponds to the geodesic matrix of distances, and for PCA, to the Euclidean distance matrix of distances. Residual errors are bound between zero and one, with zero indicating perfect reconstruction (see, e.g., Figure 2D). Note that this metric is called residual variance in the original Isomap paper^54^; we have adopted a different nomenclature to avoid confusion with our first variance explained metric.

As a third measure, we assessed the estimated dimensionality of linear and nonlinear manifolds underlying the neural population activity as function of the number of units. We posited that if a linear or nonlinear manifold captured well the geometric structure of the latent dynamics, including more units should not change the estimated dimensionality. In contrast, if the approximations provided by these methods are inaccurate, the estimated dimensionality would grow monotonically. We used the Participation Ratio^2^, PR, to estimate the dimensionality of the flat and nonlinear manifolds:

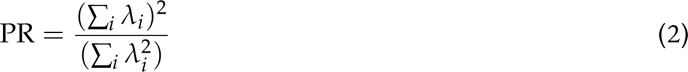

Where *λ_i_* are the eigenvalues obtained with either PCA or Isomap. Since the eigenvalues are a measure of the amount of variance explained by each mode, the Participation Ratio gives us an estimate of the variance distribution across the different modes. In the extreme case that all the eigenvalues are the same, PR equals *n*, meaning the variance is evenly distributed across all dimensions. In contrast, if PR equals one, the majority of the variance is captured by the first dimension. This way, the Participation Ratio gives an estimation of the effective dimensionality, defined as the number of dimensions necessary to explain ~ 80% of the total neural variance^2, 56^. To test the stability (or lack of) the estimated manifold dimensionality as number of neurons, we took 10 random subsets of neurons between 5 and the total number of recorded neurons, in steps of 10. Results are reported as mean±s.d.; Figure 2E shows a representative example. Note that in contrast to Ref. 2 we used single-trial activity because our goal was to consistently compare the dimensionality of manifolds for different numbers of neurons, rather than estimating the actual dimensionality of the neural population activity. Moreover, we observed that the value of the Participation Ratio did not change after enough (*∼* 15 trials per target, for the centre-out reaching task) trials were considered.

Finally, to obtain a direct comparison between the estimated dimensionality of linear and nonlinear manifolds as a function of the number of units considered for their estimation, we computed a “nonlinearity index” as the ratio between the estimated dimensionality of the linear to that of the nonlinear manifold for the same number of units (see, e.g., Figure 2F).

### Establishing the influence of task complexity on manifold nonlinearity

We investigated whether more complex behaviours requiring a broader range of actions would reveal greater manifold nonlinearity by directly comparing manifolds underlying a “simple” and a “complex” version of the same task. We defined the simple task as a subset comprising one (for the monkey data) or several conditions (for the human data) of the full set of conditions, which constituted the complex task. We matched the total number of trials between simple and complex task by randomly subsampling from the complex task while keeping the same number of trials per condition, to avoid biasing our results based on the number of data points.

#### Monkey

We defined the simple and complex tasks in the monkey data as reaching to one target and reaching eight targets, respectively. We performed the comparison between simple and complex tasks while considering each of the eight directions as a different simple task. For each reach direction, we repeated the analysis 10 times, taking different samples of trials from the complex direction. The results, shown in Figure S4E, represent the average estimated dimensionality of flat and nonlinear manifolds across all 10 repetitions of the eight simple tasks.

#### Human

For the human data, we investigated the influence of task complexity in manifold nonlinearity during one session in which the participant attempted to draw straight-line strokes, and two sessions in which the participant attempted to write single letters. For the session in which the participant attempted to draw straight-line strokes, we defined the simple and complex task as we did for the monkeys: the simple comprised lines in only one direction but considering all three lengths (for a total of 30 trials), and the complex task comprised the same number of trials including lines from all 16 directions. We repeated this analysis 10 times per reach direction (taking different subsets of trials for the complex task), and averaged the results across them.

For the sessions in which the participant attempted to draw individual letters, we define the simple and complex tasks in a slightly different way. Intuitively, and in agreement with Figure 1e in Ref. 22, similar shape letters and symbols should require more similar M1 activity patterns than letters with different shape. We thus defined the simple task by selecting five letters with similar morphology^60^. We verified that similar letters defined based on their morphology indeed required more similar activity patterns than any pair of randomly selected letters by computing the Euclidean distance between neural states (details) in the full-dimensional state space (inset in Figure 3G). For this analysis, we defined the simple task by taking 50 trials in which the participant attempted to write similar letters or symbols. As before, for the complex task, we selected a matched number of trials from all the different letter and symbols. We repeated this process for five groups of similar letters taking, for each of them, 50 subsets of trials to define the complex task. We report the average result.

### Additional analyses including controls

#### Pairwise correlation analysis

We simply defined the pairwise correlation between pairs of units as their Pearson’s correlation coefficient.

#### Simulation of neural signals

We generates synthetic data to evaluate if the nonlinearity index is representative of manifold nonlinearity. We replicated the methods in Altan et al.^55^. Initially, we generated d-dimensional signals by choosing (d x M) random samples from a distribution of firing rates derived from multi-electrode array recordings of neural activity in the macaque primary motor cortex (M1) during a center-out task. These firing rates were grouped in 30 ms intervals. The selection of samples was made randomly from all the neurons and time intervals recorded during successful tasks. These signals served as a basis of known dimension d (d=10), maintaining the initial firing statistics observed in the M1 recordings. The 10-dimensional latent signals were initially smoothed with a Gaussian kernel before being multiplied by an N × d mixing matrix W. The values of W were randomly sampled from a Gaussian distribution with a mean of zero, and a variance of one. We used N=84 to replicate the nb of neurons in the dataset used. We then created a nonlinear embedding of the sythetic neural data by processing each simulated neural recordings with a exponential activation function as descripbed bellow :

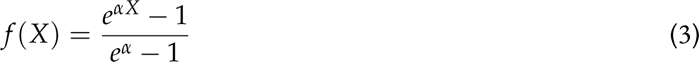

We varied the degree of nonlinearity by changing the parameter *α* between the values of 1, 4 and 8^55^.

#### Results do not depend on the specific dimensionality reduction technique used to identify the flat and nonlinear manifolds

We used PCA and Isomap as linear and nonlinear methods to estimate a flat and a nonlinear manifold underlying the neural population activity, respectively. To verify that our results were not a consequence of our choice of dimensionality reduction techniques, we repeated our main analyses using two alternative methods: Factor Analysis and Nonlinear PCA. Factor analysis (FA), like PCA, is often used to find a flat manifold underlying neural population recordings^39^. FA identifies a low-dimensional space that preserves the variance that is shared across units while discarding variance that is independent of each unit. We then estimated the dimensionality of these flat manifolds by computing the Participation Ratio using the eigenvalues of the shared covariance matrix. Importantly, the estimated dimensionality of flat manifolds identified using PCA and FA was highly correlated (Figure S3C), indicating that our results do not arise from the inherent assumptions of PCA.

Similarly, we verified that the better ability of Isomap to capture the neural population activity when compared to PCA generalised to other nonlinear dimensionality reduction techniques by repeating the main analyses using Nonlinear PCA. Nonlinear PCA is an autoencoder-based approach that finds a latent representation of the input firing rates and orders those latent signals (“nonlinear PCs”) based on their variance explained, enforcing a PCA-like structure on the nonlinear low-dimensional embedding^59^. We used the MATLAB package from Scholz et al.^59^. Since nonlinear PCA does not yield eigenvalues associated to the latent signals, explained variance was defined based on the quality of data reconstruction. We used an 80% threshold to define the dimensionality of these flat manifolds, since this is the approximate threshold provided by the Participation Ratio. Our direct comparison between nonlinear PCA and Isomap shows a high correlation between the estimated dimensionality of their respective nonlinear manifolds (Figure S3E). Thus, our results do not arise from the implicit assumptions of Isomap alone, but generalise to other nonlinear dimensionality reduction techniques.

Finally, we also verified that our results were not contingent on our choice of the Participation Ratio as a metric for manifold dimensionality estimation. Thus, we verified that our results held when considering a recently proposed principled alternative, Parallel Analysis^55^. Parallel analysis generates null distributions for the eigenvalues by repeatedly shuffling each of the *n* firing rate vectors separately. The shuffling step ensures that the remaining covariation structure across firing rates is not due to chance. Similar to Ref. 55, we repeated the shuffling procedure 200 times, resulting in a null distribution for each eigenvalue based on 200 samples. The eigenvalues that exceeded the 95*^th^* percentile of their null distribution were identified as significant; the number of significant eigenvalues determined the dimensionality of the flat manifold. Figure S3D shows that the results obtained using Parallel Analysis on the eigenvalues obtained with PCA are similar to those obtained using the Participation Ratio, establishing that the estimated dimensionality of flat manifolds does not depend on our adopting a particular metric.

#### Establishing the behavioural relevance of nonlinear manifolds

To test whether the nonlinear neural modes identified with Isomap were behaviourally relevant, we built standard Wiener filter decoders to predict continuous hand movements based on the latent dynamics within flat (identified with PCA) and nonlinear (identified with Isomap) manifolds with increasing dimensionality. For decoding, we used Ridge Regression (the Ridge class in Ref. 97). Since the hand trajectory is a two-dimensional signal, we built separate decoders to predict the hand trajectories along the X and Y axes; we reported the average performance across these two axes. The *R*^2^ value, defined as the squared correlation coefficient between actual and predicted hand trajectories, was used to quantify decoder performance. Figure S3E shows that linear decoders trained on latent dynamics within nonlinear manifolds provide more accurate behavioural predictions than their counterparts trained on latent dynamics within flat manifolds.

## Neural network models

### Recurrent neural network models

#### Model Architecture

To better understand the factors underlying manifold nonlinearity, we trained recurrent neural networks (RNNs) to perform the same centre-out reaching task as the monkeys. These models were implemented using Pytorch^98^. Similar to previous studies simulating motor cortical dynamics during reaching^18, 36, 37, 49^, we implemented the dynamical system 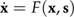 to describe the RNN dynamics:

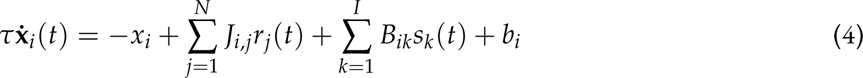

where *x_i_* is the hidden state of the *i*-th unit and *r_i_* is the corresponding firing rate following *tanh* activation of *x_i_*. All networks had *N* = 1000 units and *I* = 3 inputs, a time constant *τ* = 0.05 s, and an integration time step *dt* = 0.03 s. Each unit had an offset bias, *b_i_*, initially set to zero. The initial states *x_t_*_=0_ were sampled from the uniform random distribution (0.2, 0.2). To understand the influence of recurrent connectivity in manifold nonlinearity, we trained networks with an increasing percentage of recurrent connections (10 %, 40%, 70%, and 100 %). The recurrent weights *J_ij_* were initially sampled from the Gaussian distribution 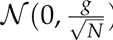, where *g* = 1.2. The time-dependent inputs *s* (specified below) fed into the network had input weights *B* initially sampled from the uniform distribution *U* (*−*0.1, 0.1).

In order to replicate experimental trial-to-trial variability across reaches to the same target, we designed the networks so they took actual preparatory activity and produced single reaches recorded during a representative session. The inputs *s* were four-dimensional, consisting of a one-dimensional fixation signal and a three-dimensional target signal. The target signal remained at 0 until the target was presented (set to *t* = 210 ms). The fixation signal started at 1 and went to 0 at movement onset (set to *t* = 420 ms). The three-dimensional target signal was derived from trial-specific preparatory activity from monkey neural data by integrating over time the latent dynamics along the first *D* = 3 neural modes obtained from performing PCA during the instructed delay period^99^ (-210 to +30 ms with respect to the go cue).

The networks were trained to produce two-dimensional outputs *v* corresponding to *x* and *y* velocities of the experimentally recorded reach trajectories, which were read out via the linear mapping:

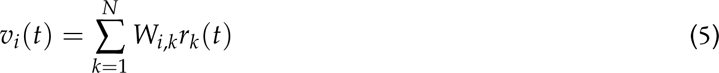

where the output weights *W* were sampled from the uniform distribution *U* (*−*0.1, 0.1).

#### Training

Networks were trained to generate velocities of experimental reach trajectories from 600 trials from Monkey C_L_ using the Adam optimiser with a learning rate *l* = 0.001, first moment estimates decay rate *β*_1_ = 0.9, second moment estimates decay rate *β*_2_ = 0.999, and *ɛ* = 1*e* 8. Networks were trained for 2500 trials with a batch size *B* = 64. The loss function *L* was defined as the mean squared error between the two-dimensional output and the target velocities over each time step *t*, with the total number of time steps *T* = 25 (equal to 750 ms with *dt* = 0.03 s):

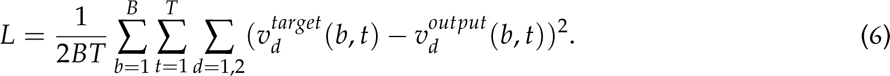

To produce dynamics that align more closely to experimentally estimated dynamics, we added L2 regularization terms for the activity rates and network weights in the overall loss function *L_R_* used for optimization^49^ :

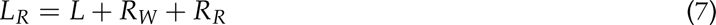

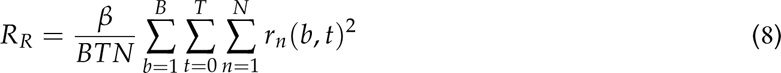

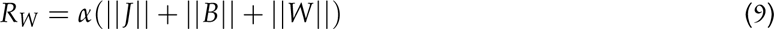

where *β* = 0.5 and *α* = 0.001. We clipped the gradient norm at 0.2 before applying the optimization step.

To model the experimental observation of lower pairwise correlations between neurons, we trained a set of “decorrelated networks” with an additional loss term that penalized the magnitude of pairwise correlations:

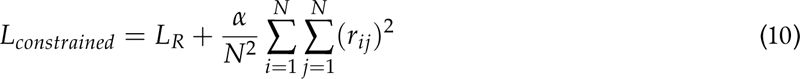

where *r_ij_* is the Pearson correlation coefficient between the *i*-th and *j*-th units, and *α* = 0.5. Both “standard” and “decorrelate” network training were performed on ten different networks initialised from different random seeds.

### LSTM model

To establish that our core model results did not depend critically on the type of neural network architecture we adopted, we repeated our main analyses using a long short-term (LSTM) architecture. We chose this architecture given its ability to predict movement during various tasks^100^. We used the same approach to replicate single trial variability as for the RNNs: we used the actual preparatory activity as inputs and trained the LSTMs on the actual hand velocities that monkeys produced during that trial.

We again trained the networks using the Adam optimiser with a learning rate *l* = 0.001, and used hyperbolic tangent (*tanh*) as the network’s nonlinearity. Networks were trained until the error, they explained at least 70% of the variance of the target velocity signals, according to the error function:

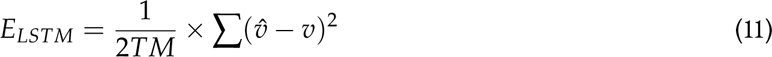

Where 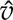 and *v* are the predicted and measured two-dimensional velocities, respectively, *T* is the number of time bins (*T* = 25), and *M* is the mini-batch size (*M* = 8 trials).

To compare the network activity and the neural data, we rectified, square-root-transformed, and smoothed the unit activity using the same parameters as for the neural recordings.

### Model validation

We used Canonical Correlation Analysis (CCA), a method that finds linear transformations that maximize the correlation between pairs of signals^101^, to quantify the similarity between the neural network activity and the neural recorded data^36, 49^. First, we separately applied PCA to the processed neural firing rates and model activations, and then performed CCA on their 15-dimensional latent dynamics; this provided a vector with 15 monotonically decreasing correlation values. To establish the relevance of these canonical correlations, we compared their values to various lower bounds obtained by shuffling the data in different ways (over time, across targets, or both over time and across targets). In all cases the actual canonical correlations greatly exceeded the shuffled lower bounds (e.g., Figure S7D, S10D).

## Data availability

Many of the monkey datasets and the human dataset used in this study are publicly available on Dryad (https://datadryad.org/stash/dataset/doi:10.5061/dryad.xd2547dktand https://datadryad.org/stash/ dataset/doi:10.5061/dryad.wh70rxwmv respectfully). The remaining data that support the findings in this study are available from the corresponding author upon reasonable request.

## Code availability

All code to reproduce the analyses in the paper will be made freely available upon peer-reviewed publication on GitHub.

## Author contributions

M.G.P. and J.A.G. devised the project. M.G.P. and L.E.M. provided the monkey datasets. J.P. and J.T.D. provided the mouse datasets. C.F. and J.B.V. analysed data, ran simulations, and generated figures. C.F., J.B.V. and J.C. provided computational models. C.F., J.B.V., J.T.D., M.G.P. and J.A.G. interpreted the data. C.F. and J.A.G. wrote the manuscript. All authors discussed and edited the manuscript.

## Acknowledgements

We thank Sara A. Solla for discussions about this research and feedback on the manuscript. L.E.M. received funding from the NIH National Institute of Neurological Disorders and Stroke (NS053603 and NS074044). J.A.G. received funding from the EPSRC (EP/T020970/1) and the European Research Council (ERC-2020-StG-949660). The funders had no role in study design, data collection and analysis, the decision to publish, or preparation of the manuscript.

## Supplementary Figures

**Figure S1:**
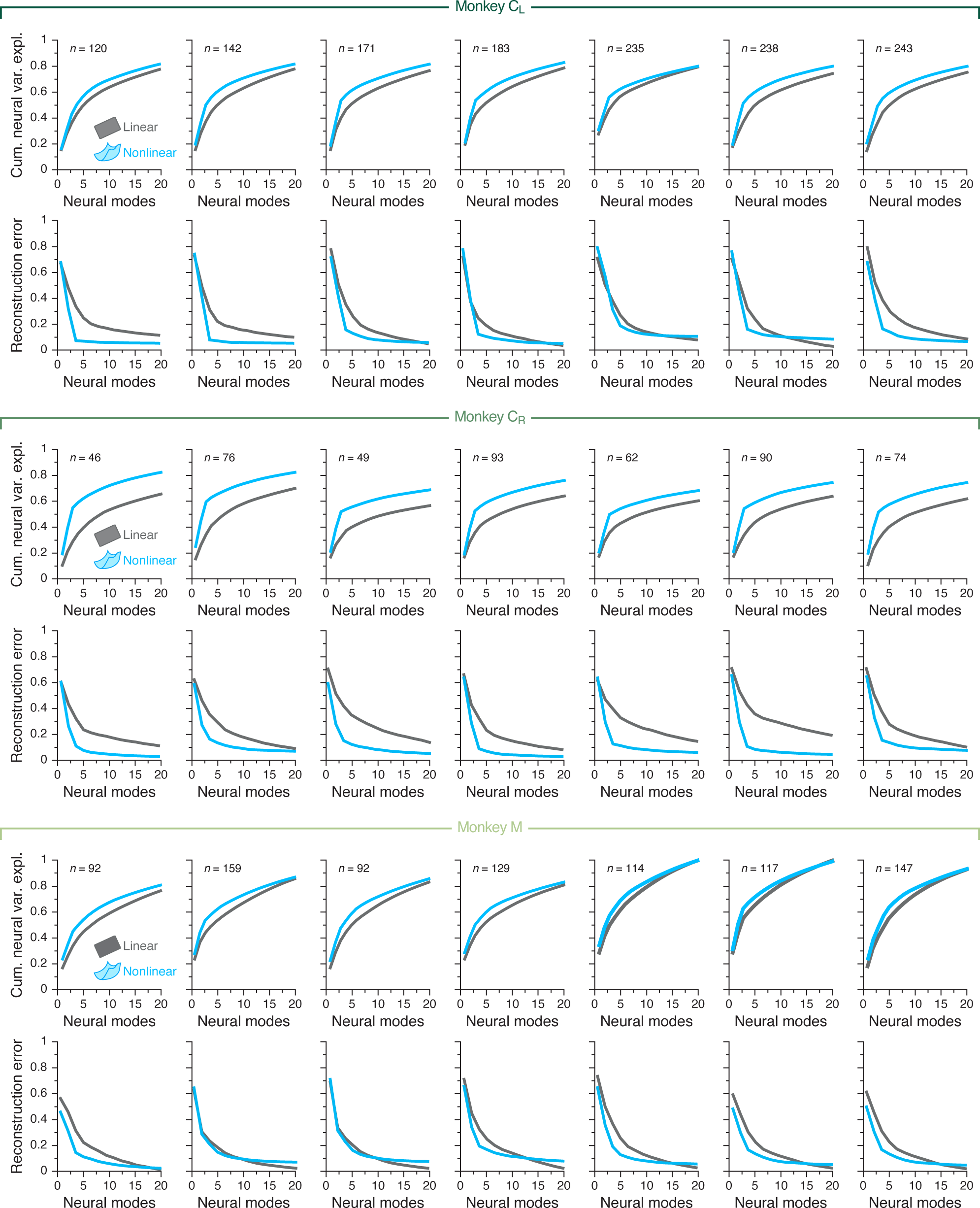
Additional data demonstrating the nonlinearity of neural manifolds underlying motor cortical population activity during a centre-out reaching task. Each pair of plots shows the cumulative neural variance explained (top) and reconstruction error (bottom) for each dataset from each monkey (header), complementing the examples in Figure 2C,D. Note that in all cases a nonlinear manifold calculated using Isomap captures the motor cortical population activity better than a flat manifold calcualted using PCA. *n*, number of recorded neurons.

**Figure S2:**
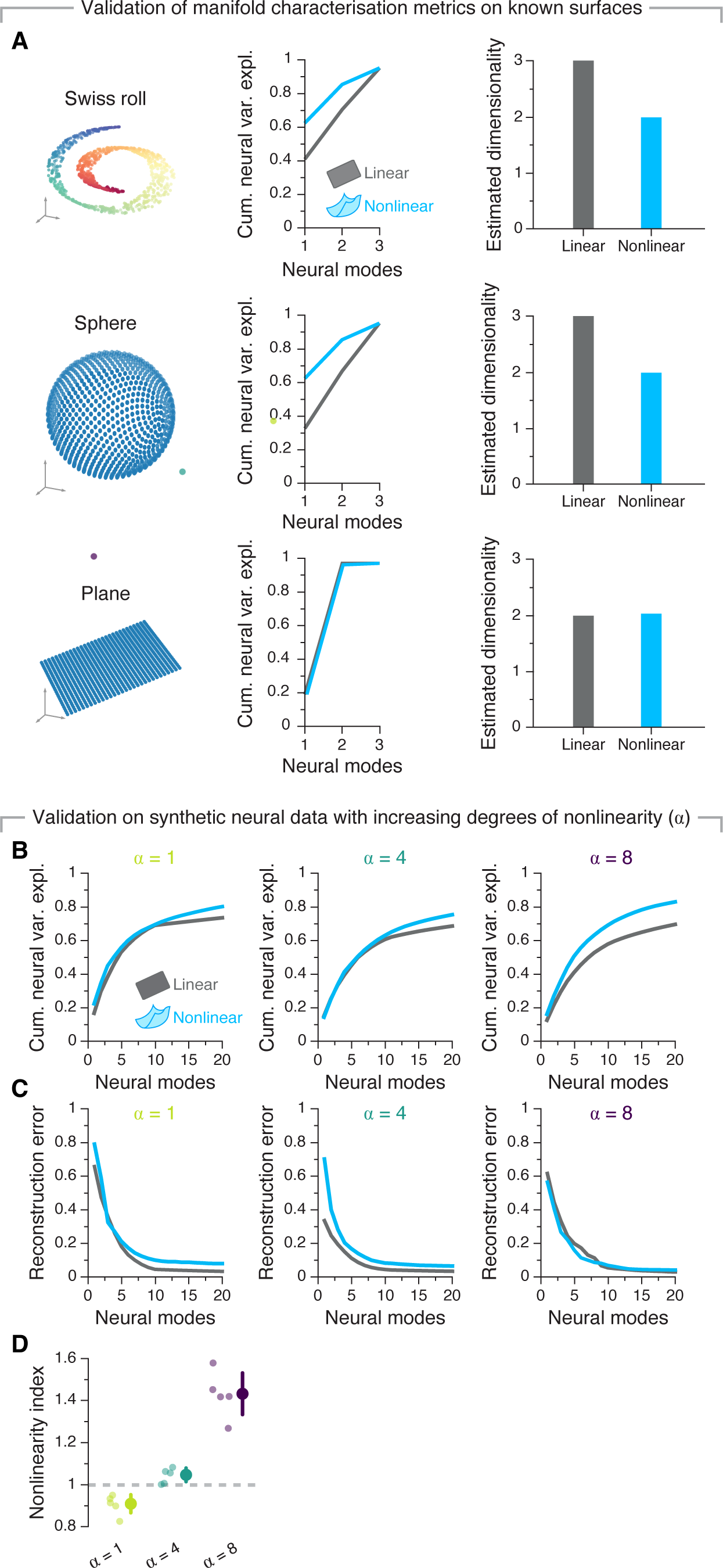
Validation of our manifold characterisation metrics on surfaces with known geometry, and synthetic neural data. **A. Left** We simulated a two-dimensional Swiss roll, a sphere and a plane embedded in a three-dimensional space. **Middle** Cumulative variance explained for each surface using PCA (grey) and Isomap (blue). Note that, while for the nonlinear case there is an elbow at two-dimensions for the Swiss roll and the sphere–indicating a potentially meaningful aspect of the data–, these are lacking in the flat case. **Right** Estimated dimensionality for the linear and nonlinear manifolds when considering all three spatial dimensions (i.e., akin to considering three neurons). As expected, Isomap provided accurate estimations, whereas PCA overestimated the dimensionality for all non-flat surfaces. We futher validated of manifold nonlinearity measures in a synthetic dataset with varying degrees of imposed nonlinearity (determined by parameter *α*; details in Methods section). **B.** Cumulative neural variance explained by flat and nonlinear manifolds for synthetic datasets with increasing degree of nonlinearity. **C.** Same as B for the reconstruction error. **D.** Nonlinearity index comparison for synthetic datasets with increasing degree of nonlinearity. Note that the index accurately captures the imposed increase in nonlinearity as the value of parameter *α* increased. Individual points represent multiple synthetic datasets generated. Marker and errorbar, mean±s.d. for each value of *α*.

**Figure S3:**
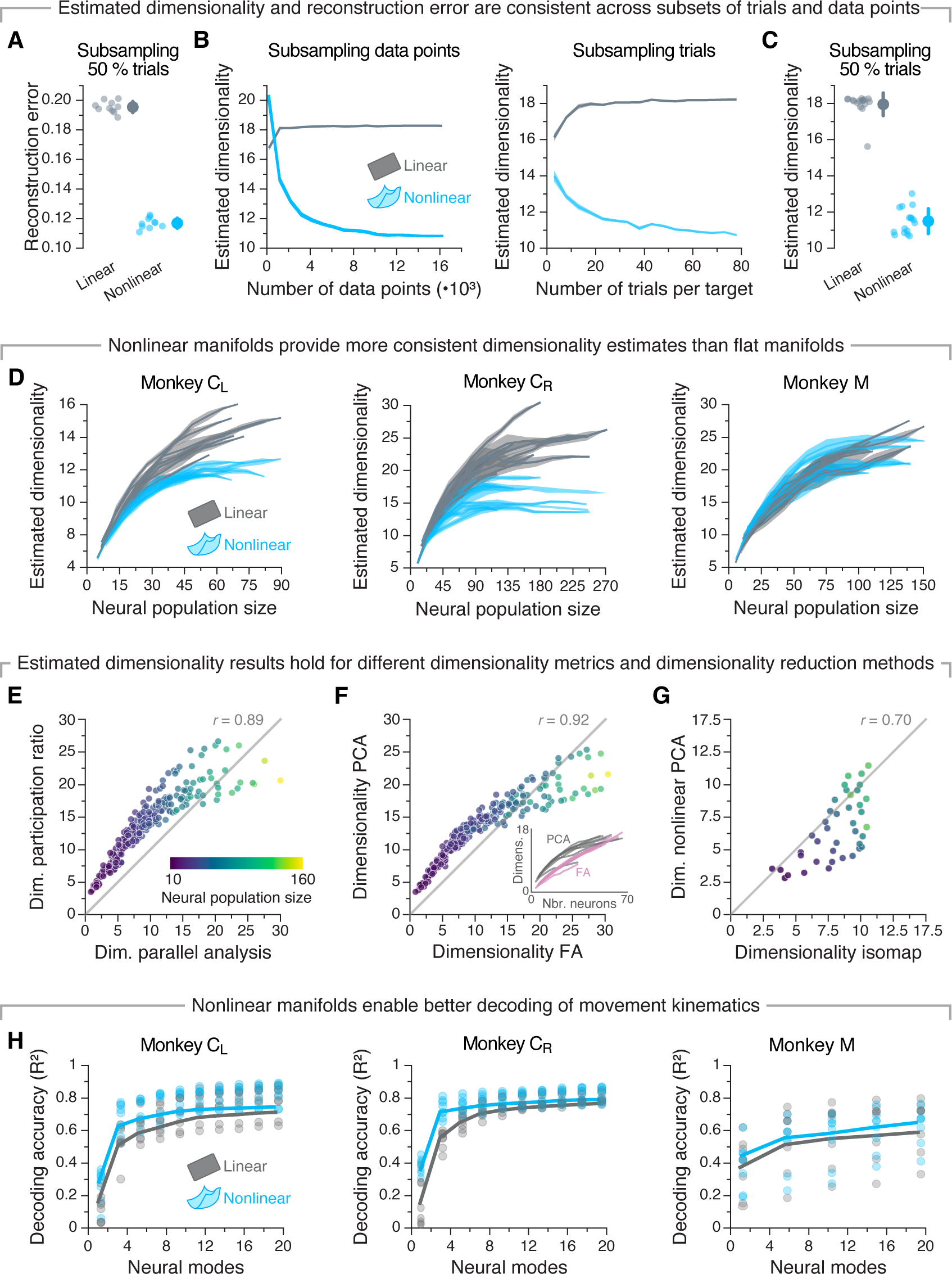
Additional data demonstrating the nonlinearity of neural manifolds underlying motor cortical population activity during a centre-out reaching task, including controls establishing the robustness of our metrics on actual neural data. **A**. Randomly subsampling trials (keeping 50% of them) does not affect the reconstruction error. Analysis done in one example session from monkey C_L_**B**. The estimated dimensionality of flat (grey) and nonlinear (light blue) manifolds plateaus after enough data points (left) or trials (right) are considered. Analysis done in one example session from monkey C_L_ **C**. Randomly subsampling trials (keeping 50% of them) does not affect the estimated dimensionality of linear or nonlinear manifolds. Analysis done in one example session from monkey C_L_**D**. Estimated dimensionality of flat (grey) and nonlinear (light blue) manifolds as function of the number of neurons used to sample them. Data from all sessions from Monkeys C_R_ (left), C_L_ (middle), and M (right); the results from Monkey C_L_ are reproduced from Fig. 2E. Mean and shaded areas, mean s.d. **E**. Comparison between the dimensionality estimated using the participation ratio and parallel analysis as function of the number of recorded neurons (legend). The results do not depend critically on the metric. Individual markers, represent one subsample of the population for all the sessions from Monkey C_L_. **F**. Comparison between the estimated dimensionality of flat manifolds computed using PCA and Factor Analysis (FA) as function of the number of recorded neurons (legend). Note the strong correlation. Data presented as in Panel E. **G**. Comparison between the estimated dimensionality of nonlinear manifolds computed using Isomap and Nonlinear PCA. Data presented as in Panel E for example session from Monkey C_L_. **H**. Accuracy of linear decoders trained to predict hand velocity as function of the number of latent signals within flat (dark blue) and nonlinear manifolds (light blue) taken as inputs. Data for all three monkey datasets. Individual markers, individual sessions; line, mean across sessions.

**Figure S4:**
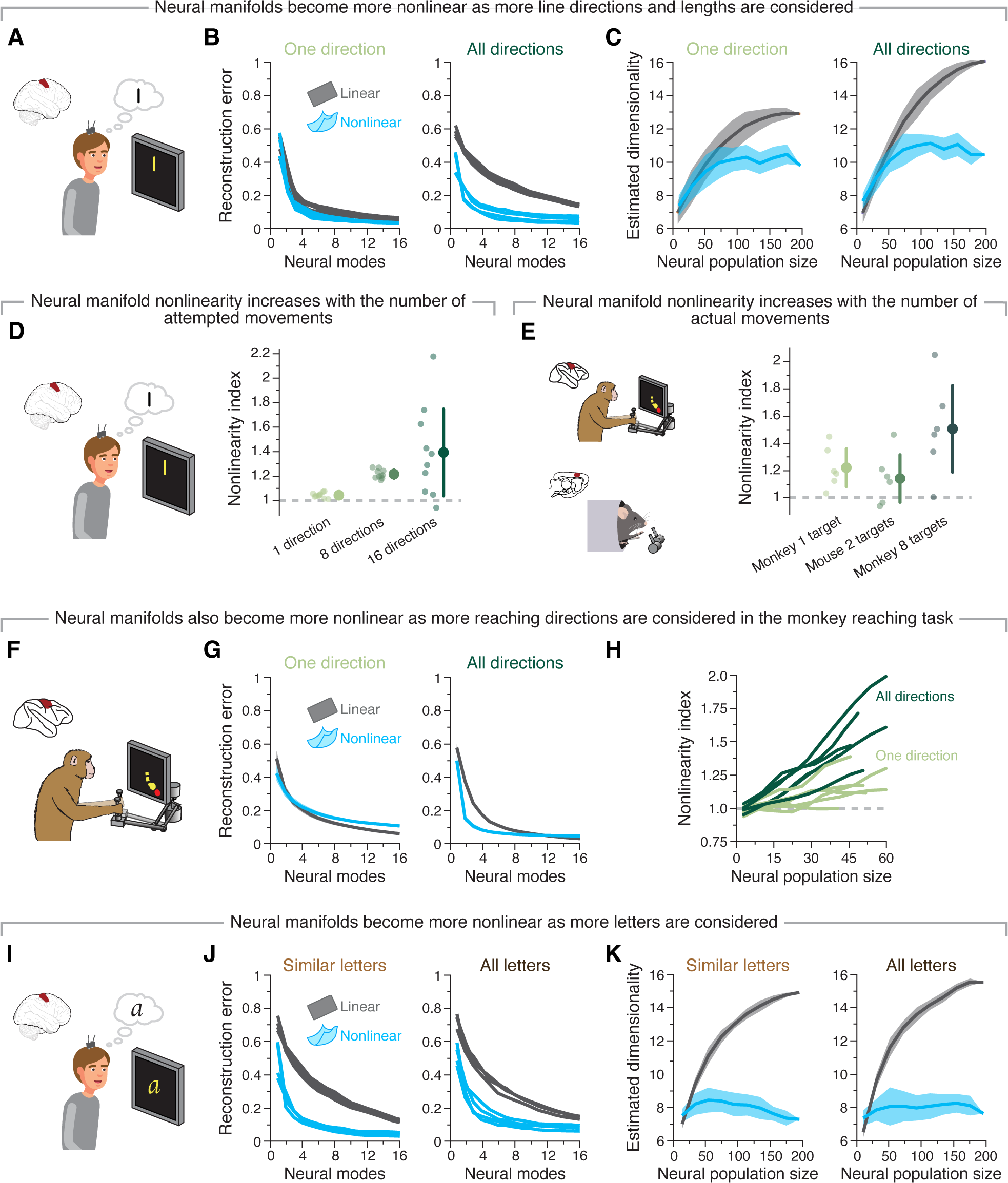
Additional data indicating that the nonlinearity of neural manifolds increases during more varied behaviours. **A**. We compared the nonlinearity underlying attempting to draw lines in one or many different directions. **B**. Reconstruction error after fitting flat (grey) and nonlinear (blue) manifolds with increasing dimensionality to neural activity recorded while attempting to draw lines in one direction (left) or all 8 directions (right). **C**. Estimated dimensionality of flat (grey) and nonlinear (blue) manifolds as function of the number of units used to sample them as the participant attempted to draw lines in one direction (left) or all 16 directions (right). Lines and shaded areas, mean. **D**. Nonlinearity index indicating the ratio of the estimated dimensionality of nonlinear manifolds to that of flat manifolds as function of the number of neurons used to sample them for different numbers of directions in the (attempted) line drawing task. Values greater than one (dashed grey line) indicate that nonlinear manifolds capture the data better than flat manifolds. Individual markers, one random selection of trials for the same dataset. Marker and errorbar, mean s.d. for each number of targets. **E**. Nonlinearity index as a function of number of targets for monkey and mouse datasets. Individual markers represent each individual session for Monkey *C_L_* and all sessions from mouse dataset. Marker and errorbar, mean s.d. for each number of targets. **F-H**. Same as A-C when comparing reaches to one vs. all eight directions for the monkey centre-out reaching task. **I-K**. Same as A-C but for the (attempted) writing task. Similar letters were chosen as in Figure 3.

**Figure S5:**
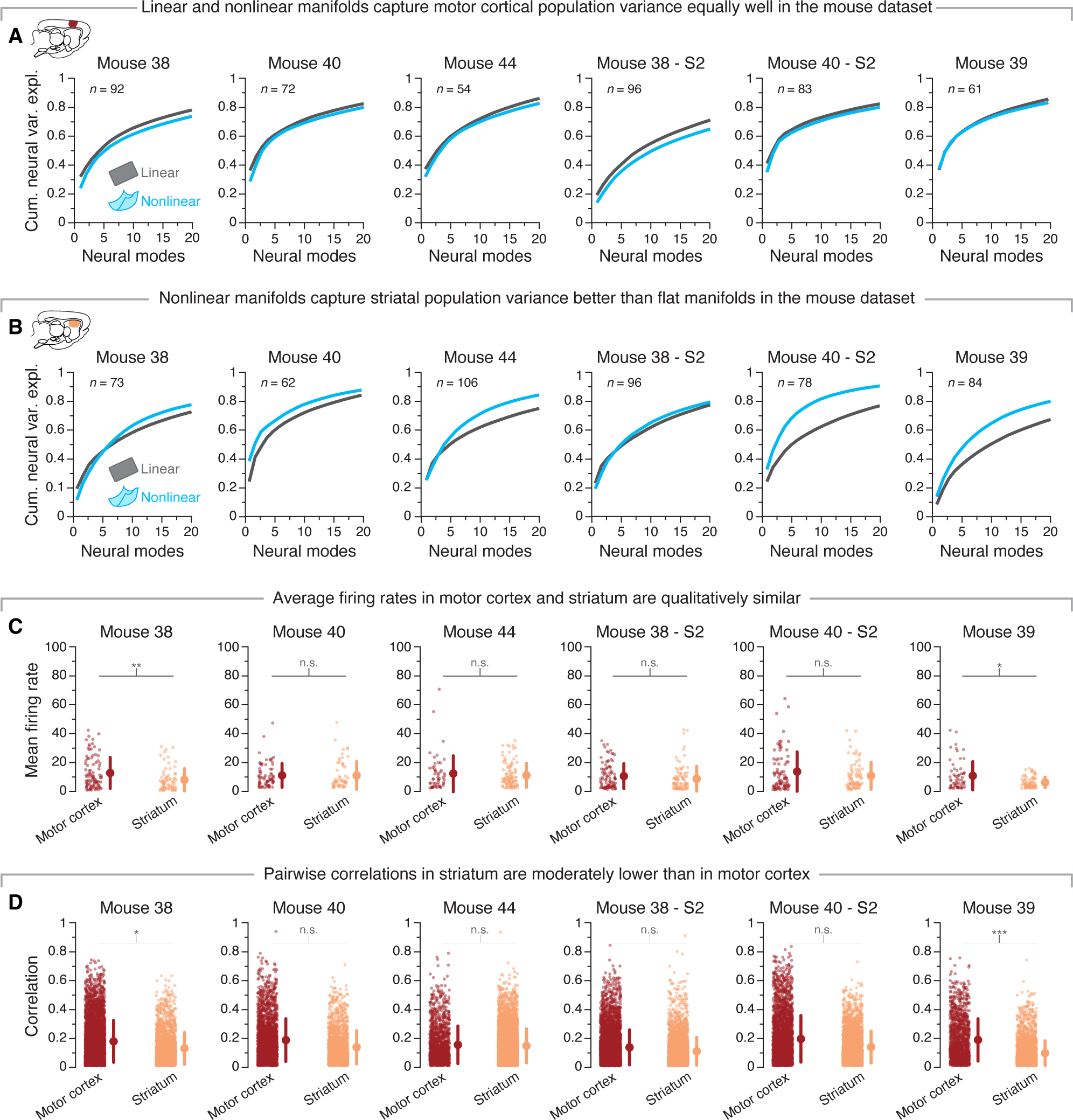
Additional data investigating the differences in the nonlinearity of motor cortical and striatal manifolds in mice engaged in a reaching, grasping, and pulling task. **A**. Cumulative neural variance explained by flat (grey) and nonlinear (blue) motor cortical manifolds as function of the number dimensions for all six mouse datasets. **B**. Cumulative neural variance explained by flat (grey) and nonlinear (blue) striatal manifolds as function of the number dimensions for all six mouse datasets. **C**. Comparison of single neuron mean firing rate between motor cortex and striatum. Individual markers, single neurons; marker and error bar, mean s.d. for each region. Note the similarity between regions. * 0.01 *P <* 0.05, ** 0.001 *P <* 0.01, two-sided Wilcoxon rank-sum test. **D**. Comparison of pairwise correlation strength between motor cortical (left) and striatal (right) neurons. Individual markers, Strength of pairwise correlations between one pair of motor cortical (left) or striatal (right) neurons. Marker and error bar, mean s.d. for each region. Note the moderate difference between regions. * 0.01 *P <* 0.05, ** 0.001 *P <* 0.01, two-sided Wilcoxon rank-sum test.

**Figure S6:**
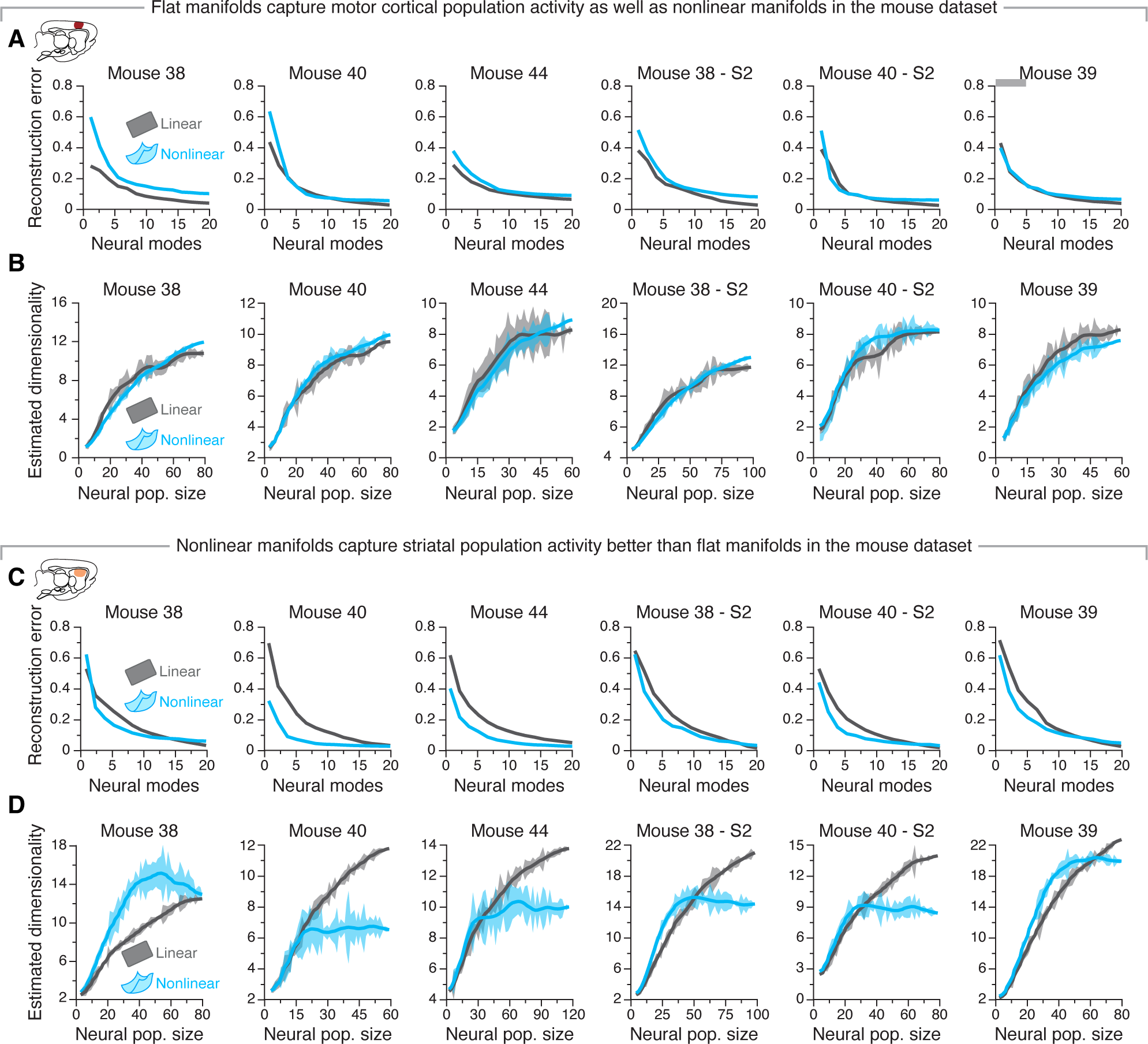
Additional data indicating differences in the nonlinearity of motor cortical and striatal manifolds in mice engaged in a reaching, grasping, and pulling task. **A**. Reconstruction error after fitting flat (grey) and nonlinear (blue) manifolds with increasing dimensionality to all the motor cortical datasets. **B**. Reconstruction error after fitting flat (grey) and nonlinear (blue) manifolds with increasing dimensionality to all the motor cortical datasets. Note that, in agreement to the reconstruction error results shown in A, flat and nonlinear manifolds seem to capture motor cortical activity equally well. **C**. Estimated dimensionality of flat (grey) and nonlinear (blue) manifolds as function of the number of neurons used to sample them. Lines and shaded areas, mean s.d. for each session. **D**. Estimated dimensionality of flat (grey) and nonlinear (blue) manifolds as function of the number of neurons used to sample them. Lines and shaded areas, mean s.d. for each session. Note the striking contrast between these results and their motor cortical counterparts (Panel B).

**Figure S7:**
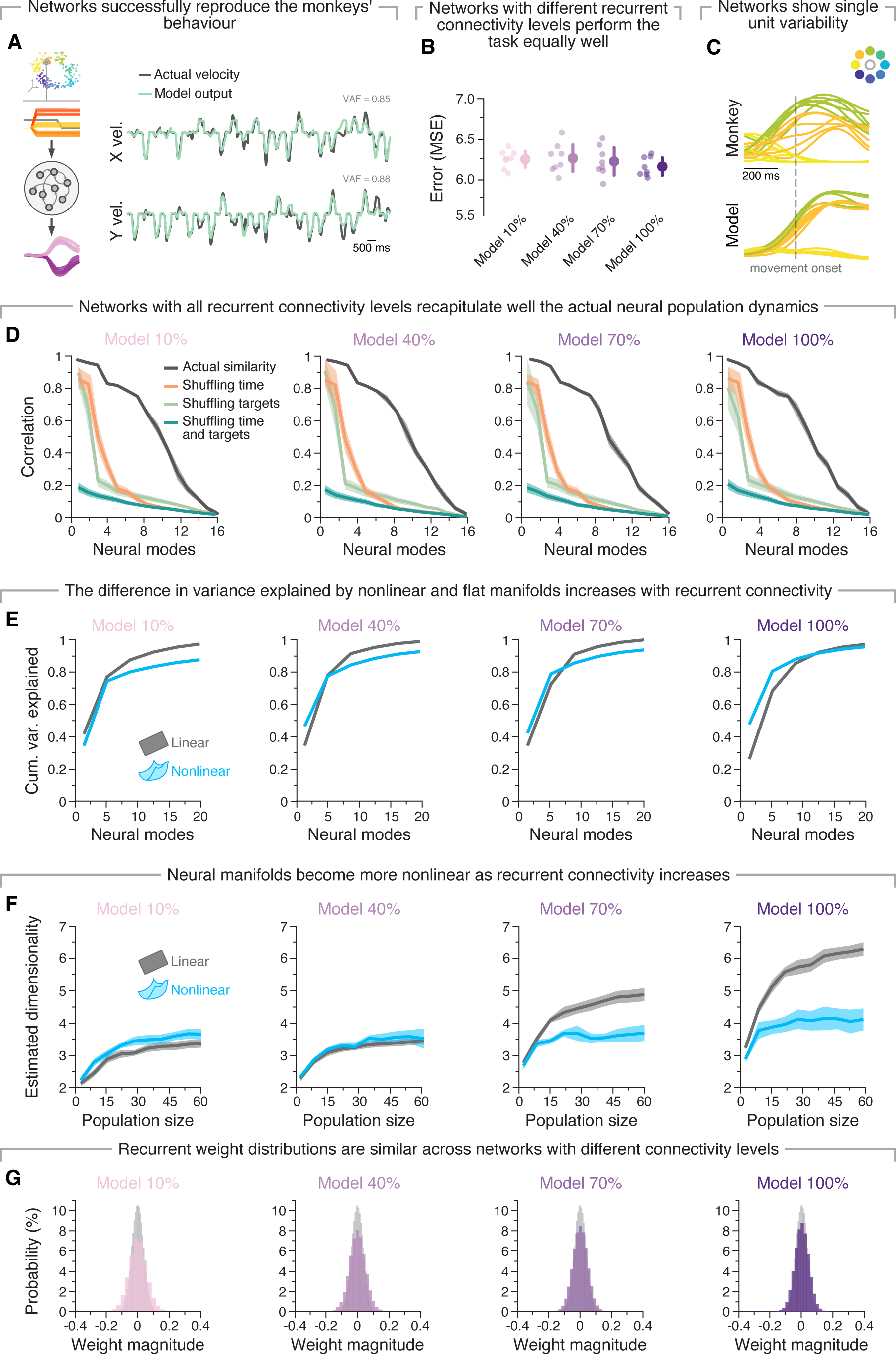
Additional data establishing the similarity of our recurrent neural network model of single reaches and actual motor cortical activity from monkeys performing the same task. **A**. Produced and target velocities along the X (top) and Y (bottom) axes for several concatenated trials. **B**. Task performance is similar across **models** with different degrees of recurrent connectivity. Individual markers, individual seeds; marker and error bars, mean s.d. across seeds. **C**. As in monkey motor cortex (top), model units (bottom) exhibit variability across trials to the same target, albeit of smaller magnitude. **D**. Similarity between the actual and simulated latent dynamics (red) for models with different degrees of recurrent connectivity, quantified using canonical correlation analysis. Note that the strength of the correlations between model and actual data is similar across different connectivity levels, whereas shuffling the model data over time (orange), across targets (light green), or across both time and targets (teal) greatly reduces this similarity. Line and shaded area, mean s.d. across seeds. **E**. Cumulative network variance explained by flat (grey) and nonlinear (blue) manifolds as function of the number dimensions for models with increasing levels of recurrent connectivity (10%, 40%, 70% and 100%). **F**. Estimated dimensionality of flat (grey) and nonlinear (blue) manifolds as function of the number of units used to sample them for models with increasing levels of recurrent connectivity (10%, 40%, 70% and 100%). Lines and shaded areas, mean s.d. across 10 random subsets of units. **G**. Connectivity matrix weight distribution before (light grey) and after training (colour) for models with increasing levels of recurrent connectivity (10%, 40%, 70% and 100%).

**Figure S8:**
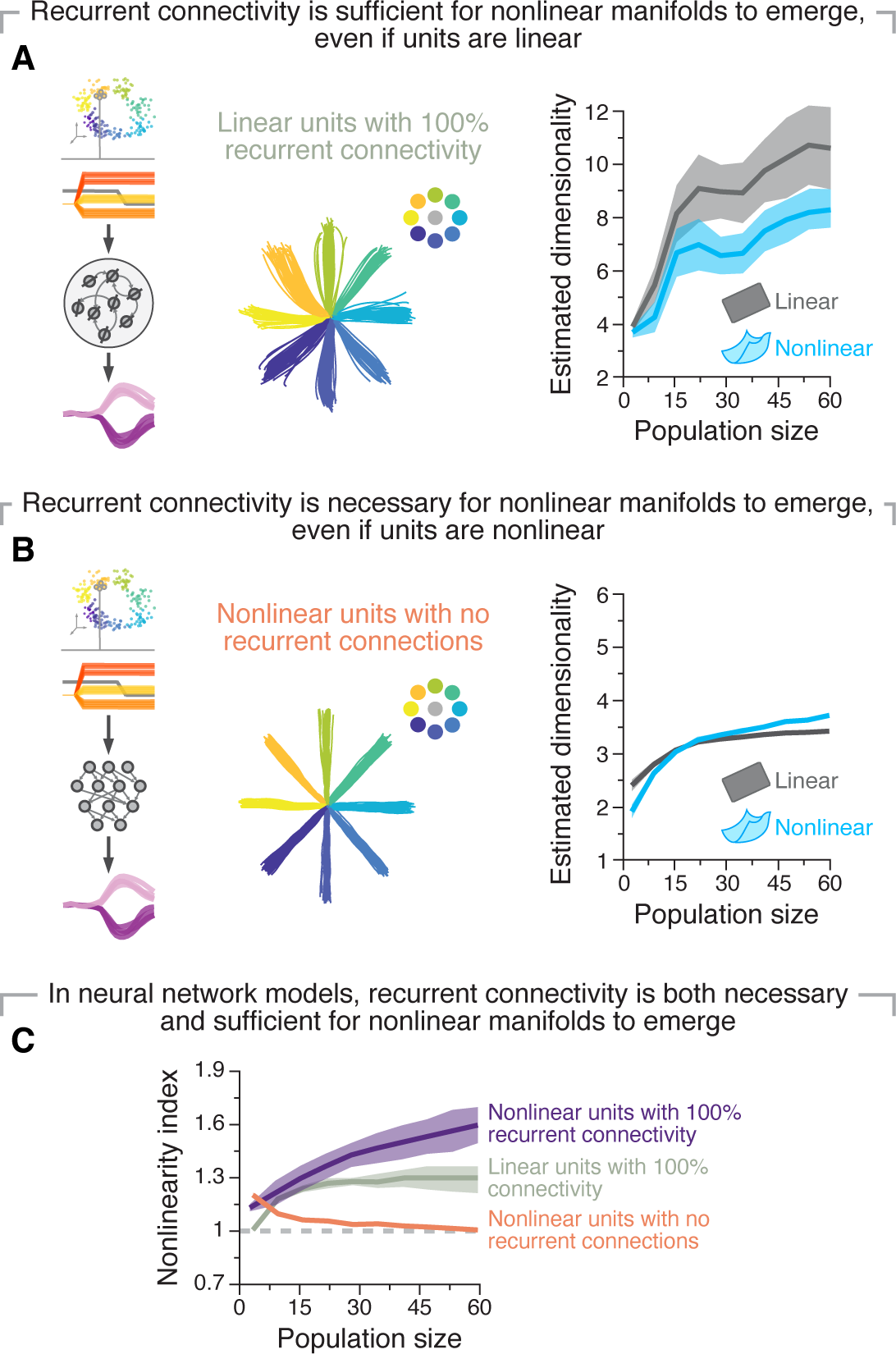
Dense recurrent connectivity is both necessary and sufficient in order for manifolds to be nonlinear in neural network models. **A**. RNNs with linear units (left) trained on the monkey centre-out reaching task (middle, example outputs) produced activity whose properties are more consistently estimated with nonlinear manifolds, even if the units are linear(right). Line and shaded areas, mean s.d. across seeds. **B**. Fully connected feedforward networks with nonlinear units (left) trained on the monkey centre-out reaching task (middle, example outputs) produced activity whose properties are equally well estimated with flat and nonlinear manifolds (right). Data presented as in A. **C**. Nonlinearity index indicating the ratio of the estimated dimensionality of nonlinear manifolds to that of flat manifolds as function of the number of sampled units for the network architectures in A and B as well as the standard recurrent networks used in the paper (e.g., in Fig. 5). Combined, these results indicate that in neural network models, dense recurrent connectivity is both necessary and sufficient in order for manifolds to be nonlinear. Lines and shaded areas, mean*±*s.d. across 10 random subsets of units.

**Figure S9:**
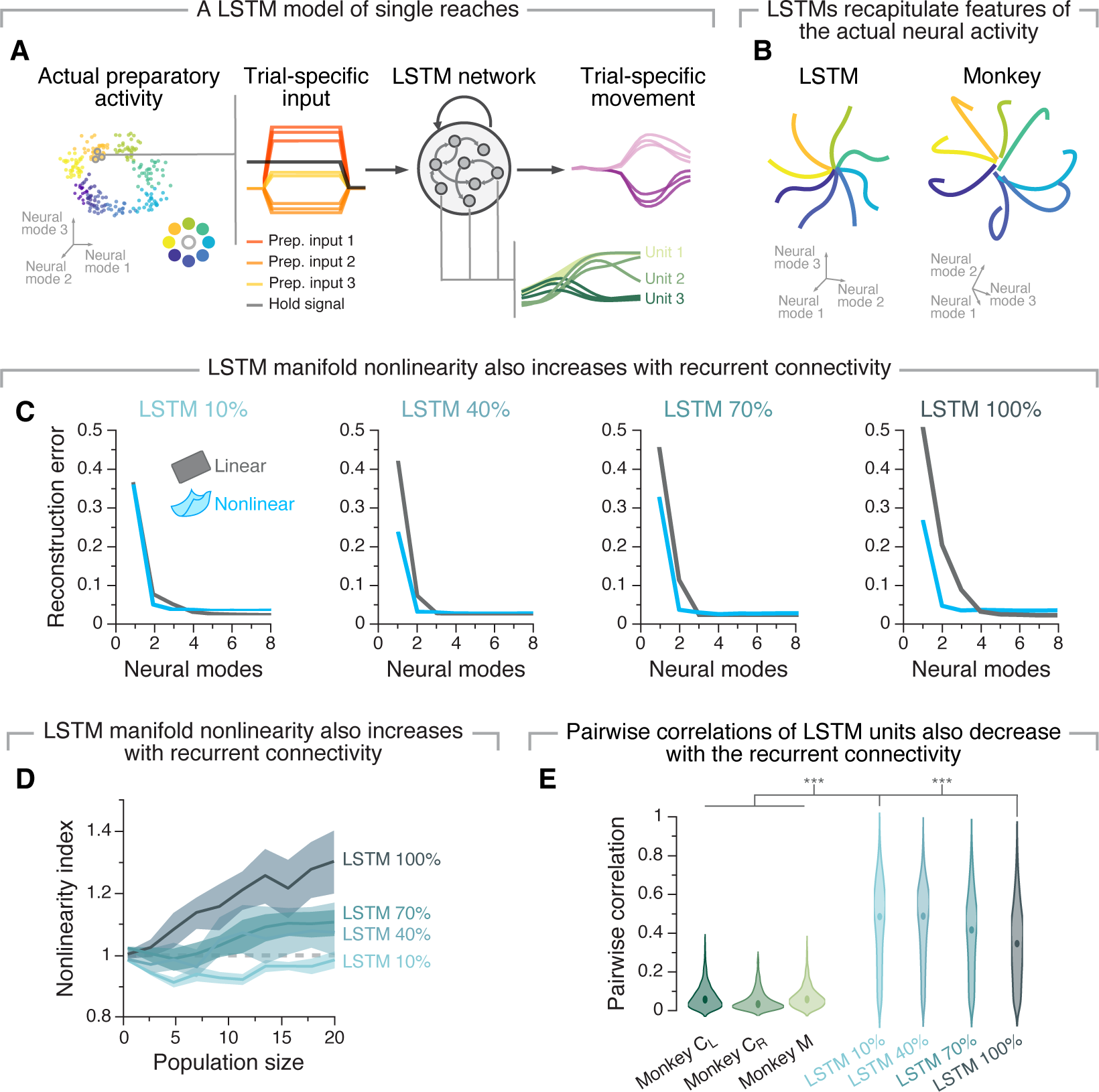
A different neural network architecture further establishes that dense recurrent connectivity is necessary for the emergence of nonlinear manifolds. **A**. We trained LSTM models to produce the hand velocities generated by the monkeys using trial-specific preparatory as inputs, replicating the training procedure for RNNs. Bottom right: the activity of three randomly selected neurons showcases variability across trials to the same target. **B**. The produced network activity recapitulated key features of neural population activity, and also exhibited single unit variability across reaches to the same target (example in Panel A, inset). **C**. Reconstruction error after fitting flat (grey) and nonlinear (blue) manifolds with increasing dimensionality. Note that the relationship between reconstruction error and network connectivity replicates that observed for RNN models (Figure 5D). **D.** Nonlinearity index indicating the ratio of the estimated dimensionality of nonlinear manifolds to that of flat manifolds as function of the number of units used to sample them for different levels of network connectivity (legend). Line and shaded area, mean s.d. across 10 seeds. **E.** The strength of pairwise correlations between network units decreases with increasing network connectivity. Shown are the strength of the pairwise correlations across all recorded neurons for all each of the three monkeys (left) and for networks with increasing connectivity levels (right). *** *P <* 0.001, two-sided Wilcoxon rank-sum test.

**Figure S10:**
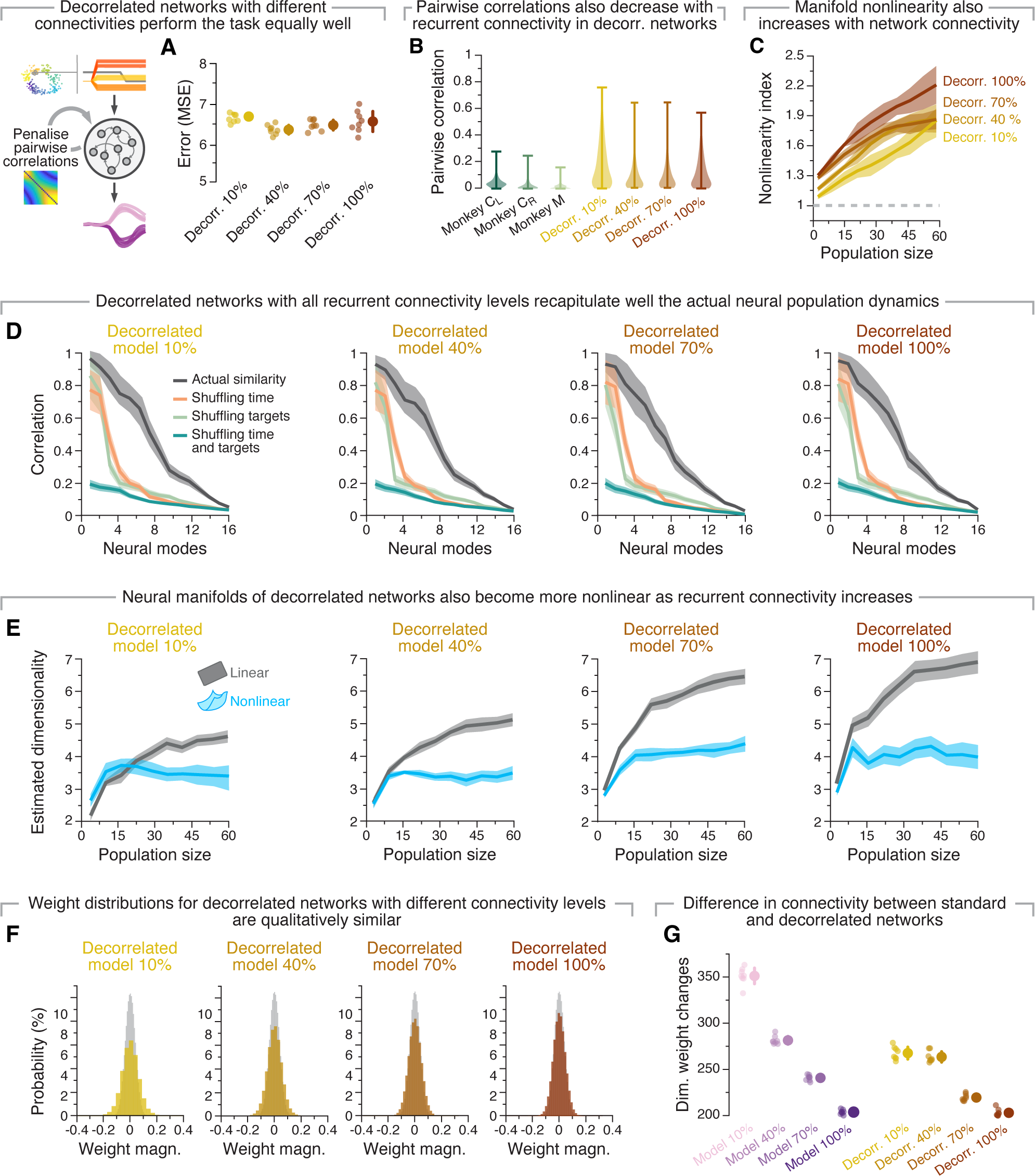
Additional data on our recurrent neural network model constrained to have reduced pairwise correlations between units. **A**. Task performance is similar across models with different degrees of recurrent connectivity. Individual markers, individual seeds; marker and error bars, mean s.d. across seeds. **B**. Strength of pairwise correlations between units from “standard networks” and “decorrelated networks” with four degrees of recurrent connectivity compared. Violin, probability density for each network architecture and recurrent connectivity level. **C**. Nonlinearity index indicating the ratio of the estimated dimensionality of nonlinear manifolds to that of flat manifolds as function of the number of sampled units for decorrelated networks with varying degrees of recurrent connectivity (10%, 40%, 70% and 100%). Lines and shaded areas, mean s.d. across seeds. Note that, in contrast to “standard networks” (5), even sparsely connected networks are now associated with clearly nonlinear manifolds. **D**. Similarity between the actual and simulated latent dynamics (red) for models with different degrees of recurrent connectivity, quantified using canonical correlation analysis. Note that the strength of the correlations between the decorrelated models and the actual data is similar across different connectivity levels, whereas shuffling the model data over time (orange), across targets (light green), or across both time and targets (teal) greatly reduces this similarity. Line and shaded area, mean s.d. across seeds. **E**. Estimated dimensionality of flat (grey) and nonlinear (blue) manifolds as a function of the number of units used to sample them for decorrelated networks with increasing levels of recurrent connectivity (10%, 40%, 70% and 100%). Lines and shaded areas, mean s.d. across 10 random subsets of units. **F**. Connectivity matrix weight distribution before training (light grey) and after training (colour) for decorrelated models with increasing levels of recurrent connectivity (10%, 40%, 70% and 100%) **G**. A comparison of the dimensionality of the weight changes across “standard” and “decorrelated” networks with different recurrent connectivity levels following training identifies changes in network connectivity structure. **G**. A comparison of the dimensionality of the weight changes across “standard” and “decorrelated” networks with different recurrent connectivity levels following training identifies changes in network connectivity structure.

## References

1. Juan A. Gallego, Matthew G. Perich, Lee E. Miller, and Sara A. Solla. Neural manifolds for the control of movement. Neuron, 94(5):978–984, 2017.

2. Peiran Gao, Eric Trautmann, Byron Yu, Gopal Santhanam, Stephen Ryu, Krishna Shenoy, and Surya Ganguli. A theory of multineuronal dimensionality, dynamics and measurement. BioRxiv, page 214262, 2017.

3. Mark Stopfer, Vivek Jayaraman, and Gilles Laurent. Intensity versus identity coding in an olfactory system. Neuron, 39(6):991–1004, 2003.

4. Juri Minxha, Ralph Adolphs, Stefano Fusi, Adam N Mamelak, and Ueli Rutishauser. Flexible recruitment of memory-based choice representations by the human medial frontal cortex. Science, 368(6498), 2020.

5. Patrick T Sadtler, Kristin M Quick, Matthew D Golub, Steven M Chase, Stephen I Ryu, Elizabeth C Tyler-Kabara, M Yu Byron, and Aaron P Batista. Neural constraints on learning. Nature, 512(7515):423–426, 2014.

6. James H Marshel, Yoon Seok Kim, Timothy A Machado, Sean Quirin, Brandon Benson, Jonathan Kadmon, Cephra Raja, Adelaida Chibukhchyan, Charu Ramakrishnan, Masatoshi Inoue, et al. Cortical layer–specific critical dynamics triggering perception. Science, 365(6453):eaaw5202, 2019.

7. Richard J Gardner, Erik Hermansen, Marius Pachitariu, Yoram Burak, Nils A Baas, Benjamin A Dunn, May-Britt Moser, and Edvard I Moser. Toroidal topology of population activity in grid cells. Nature, 602(7895):123–128, 2022.

8. Rishidev Chaudhuri, Berk Gerç ek, Biraj Pandey, Adrien Peyrache, and Ila Fiete. The intrinsic attractor manifold and population dynamics of a canonical cognitive circuit across waking and sleep. Nature neuroscience, 22(9):1512–1520, 2019.

9. Benjamin R Cowley, Adam C Snyder, Katerina Acar, Ryan C Williamson, M Yu Byron, and Matthew A Smith. Slow drift of neural activity as a signature of impulsivity in macaque visual and prefrontal cortex. Neuron, 108(3):551–567, 2020.

10. Jay A Hennig, Emily R Oby, Matthew D Golub, Lindsay A Bahureksa, Patrick T Sadtler, Kristin M Quick, Stephen I Ryu, Elizabeth C Tyler-Kabara, Aaron P Batista, Steven M Chase, et al. Learning is shaped by abrupt changes in neural engagement. Nature Neuroscience, 24(5):727–736, 2021.

11. Juan A Gallego, Matthew G Perich, Stephanie N Naufel, Christian Ethier, Sara A Solla, and Lee E Miller. Cortical population activity within a preserved neural manifold underlies multiple motor behaviors. Nature communications, 9(1):1–13, 2018.

12. Andrew Miri, Claire L Warriner, Jeffrey S Seely, Gamaleldin F Elsayed, John P Cunningham, Mark M Churchland, and Thomas M Jessell. Behaviorally selective engagement of short-latency effector pathways by motor cortex. Neuron, 95(3):683–696, 2017.

13. Juan A Gallego, Matthew G Perich, Raeed H Chowdhury, Sara A Solla, and Lee E Miller. Long-term stability of cortical population dynamics underlying consistent behavior. Nature neuroscience, 23(2):260–270, 2020.

14. Mostafa Safaie, Joanna C Chang, Junchol Park, Lee E Miller, Joshua T Dudman, Matthew G Perich, and Juan A Gallego. Preserved neural dynamics across animals performing similar behaviour. Nature, 623(7988):765–771, 2023.

15. Emily R Oby, Matthew D Golub, Jay A Hennig, Alan D Degenhart, Elizabeth C Tyler-Kabara, M Yu Byron, Steven M Chase, and Aaron P Batista. New neural activity patterns emerge with long-term learning. Proceedings of the National Academy of Sciences, 116(30):15210–15215, 2019.

16. Matthew G Perich, Juan A Gallego, and Lee E Miller. A neural population mechanism for rapid learning. Neuron, 100(4):964–976, 2018.

17. Xulu Sun, Daniel J O’Shea, Matthew D Golub, Eric M Trautmann, Saurabh Vyas, Stephen I Ryu, and Krishna V Shenoy. Cortical preparatory activity indexes learned motor memories. Nature, 602(7896):274–279, 2022.

18. Barbara Feulner and Claudia Clopath. Neural manifold under plasticity in a goal driven learning behaviour. PLoS computational biology, 17(2):e1008621, 2021.

19. Saurabh Vyas, Nir Even-Chen, Sergey D. Stavisky, Stephen I. Ryu, Paul Nuyujukian, and Krishna V. Shenoy. Neural Population Dynamics Underlying Motor Learning Transfer. Neuron, 97(5):1177–1186.e3, March 2018.

20. Matthew T Kaufman, Mark M Churchland, Stephen I Ryu, and Krishna V Shenoy. Cortical activity in the null space: permitting preparation without movement. Nature neuroscience, 17(3):440–448, 2014.

21. Mark M Churchland, John P Cunningham, Matthew T Kaufman, Stephen I Ryu, and Krishna V Shenoy. Cortical preparatory activity: representation of movement or first cog in a dynamical machine? Neuron, 68(3):387–400, 2010.

22. Francis R Willett, Donald T Avansino, Leigh R Hochberg, Jaimie M Henderson, and Krishna V Shenoy. Highperformance brain-to-text communication via handwriting. Nature, 593(7858):249–254, 2021.

23. Britton A Sauerbrei, Jian-Zhong Guo, Jeremy D Cohen, Matteo Mischiati, Wendy Guo, Mayank Kabra, Nakul Verma, Brett Mensh, Kristin Branson, and Adam W Hantman. Cortical pattern generation during dexterous movement is input-driven. Nature, 577(7790):386–391, 2020.

24. Valerio Mante, David Sussillo, Krishna V. Shenoy, and William T. Newsome. Context-dependent computation by recurrent dynamics in prefrontal cortex. Nature, 503(7474):78–84, November 2013.

25. Christian K. Machens, Ranulfo Romo, and Carlos D. Brody. Functional, But Not Anatomical, Separation of “What” and “When” in Prefrontal Cortex. Journal of Neuroscience, 30(1):350–360, January 2010.

26. K. L. Briggman, H. D. I. Abarbanel, and W. B. Kristan. Optical imaging of neuronal populations during decisionmaking. Science, 307(5711):896–901, 2005.

27. Lara M. Boyle, Lorenzo Posani, Sarah Irfan, Steven A. Siegelbaum, and Stefano Fusi. Tuned geometries of hippocampal representations meet the computational demands of social memory. Neuron, 2024.

28. Evan D Remington, Seth W Egger, Devika Narain, Jing Wang, and Mehrdad Jazayeri. A dynamical systems perspective on flexible motor timing. Trends in cognitive sciences, 22(10):938–952, 2018.

29. Jing Wang, Devika Narain, Eghbal A Hosseini, and Mehrdad Jazayeri. Flexible timing by temporal scaling of cortical responses. Nature neuroscience, 21(1):102–110, 2018.

30. Tiago Monteiro, Filipe S Rodrigues, Margarida Pexirra, Bruno F Cruz, Ana I Gonç alves, Pavel E Rueda-Orozco, and Joseph J Paton. Using temperature to analyse the neural basis of a latent temporal decision. bioRxiv, pages 2020–08, 2020.

31. Edward H Nieh, Manuel Schottdorf, Nicolas W Freeman, Ryan J Low, Sam Lewallen, Sue Ann Koay, Lucas Pinto, Jeffrey L Gauthier, Carlos D Brody, and David W Tank. Geometry of abstract learned knowledge in the hippocampus. Nature, 595(7865):80–84, 2021.

32. Carsen Stringer, Marius Pachitariu, Nicholas Steinmetz, Matteo Carandini, and Kenneth D Harris. Highdimensional geometry of population responses in visual cortex. Nature, 571(7765):361–365, 2019.

33. Ramon Nogueira, Chris C. Rodgers, Randy M. Bruno, and Stefano Fusi. The geometry of cortical representations of touch in rodents. Nature Neuroscience, pages 1–12, January 2023. Publisher: Nature Publishing Group.

34. David Dahmen, Moritz Layer, Lukas Deutz, Paulina Anna Dabrowska, Nicole Voges, Michael von Papen, Thomas Brochier, Alexa Riehle, Markus Diesmann, Sonja Grü n, and Moritz Helias. Global organization of neuronal activity only requires unstructured local connectivity. eLife, 11:e68422, 2022.

35. SueYeon Chung and Larry F Abbott. Neural population geometry: An approach for understanding biological and artificial neural networks. Current opinion in neurobiology, 70:137–144, 2021.

36. Barbara Feulner, Matthew G Perich, Raeed H Chowdhury, Lee E Miller, Juan A Gallego, and Claudia Clopath. Small, correlated changes in synaptic connectivity may facilitate rapid motor learning. Nature communications, 13(1):1–14, 2022.

37. Joanna C. Chang, Matthew G. Perich, Lee E. Miller, Juan A. Gallego, and Claudia Clopath. De novo motor learning creates structure in neural activity space that shapes adaptation, May 2023.

38. Alexandre Payeur, Amy L Orsborn, and Guillaume Lajoie. Neural manifolds and learning regimes in neural-interface tasks. bioRxiv, pages 2023–03, 2023.

39. John P Cunningham and M Yu Byron. Dimensionality reduction for large-scale neural recordings. Nature neuroscience, 17(11):1500–1509, 2014.

40. Rufus Mitchell-Heggs, Seigfred Prado, Giuseppe P. Gava, Mary Ann Go, and Simon R. Schultz. Neural manifold analysis of brain circuit dynamics in health and disease. Journal of Computational Neuroscience, 51(1):1–21, 2023.

41. Christopher Langdon, Mikhail Genkin, and Tatiana A. Engel. A unifying perspective on neural manifolds and circuits for cognition. Nature Reviews Neuroscience, pages 1–15, April 2023.

42. Gouki Okazawa, Christina E Hatch, Allan Mancoo, Christian K Machens, and Roozbeh Kiani. Representational geometry of perceptual decisions in the monkey parietal cortex. Cell, 184(14):3748–3761, 2021.

43. Katarzyna Jurewicz, Brianna J Sleezer, Priyanka S Mehta, Benjamin Y Hayden, and R Becket Ebitz. Irrational choices via a curvilinear representational geometry for value. bioRxiv, 2022.

44. Valentino Braitenberg and Almut Schūz. Cortex: statistics and geometry of neuronal connectivity. Springer Science & Business Media, 2013.

45. Kenneth D Harris and Gordon MG Shepherd. The neocortical circuit: themes and variations. Nature neuroscience, 18(2):170–181, 2015.

46. Szabolcs Horvát, Răzvan Gămănut, Mária Ercsey-Ravasz, Lūıc Magrou, Bianca Gămănut, David C Van Essen, Andreas Burkhalter, Kenneth Knoblauch, Zoltán Toroczkai, and Henry Kennedy. Spatial embedding and wiring cost constrain the functional layout of the cortical network of rodents and primates. PLoS biology, 14(7):e1002512, 2016.

47. Junchol Park, Peter Polidoro, Catia Fortunato, Jon A Arnold, Brett D Mensh, Juan Alvaro Gallego, and Joshua T Dudman. Conjoint specification of action by neocortex and striatum. bioRxiv, pages 2023–10, 2023.

48. Nicholas A Steinmetz, Cagatay Aydin, Anna Lebedeva, Michael Okun, Marius Pachitariu, Marius Bauza, Maxime Beau, Jai Bhagat, Claudia Bōhm, Martijn Broux, et al. Neuropixels 2.0: A miniaturized high-density probe for stable, long-term brain recordings. Science, 372(6539):eabf4588, 2021.

49. David Sussillo, Mark M Churchland, Matthew T Kaufman, and Krishna V Shenoy. A neural network that finds a naturalistic solution for the production of muscle activity. Nature neuroscience, 18(7):1025–1033, 2015.

50. Jonathan A Michaels, Benjamin Dann, and Hansjōrg Scherberger. Neural population dynamics during reaching are better explained by a dynamical system than representational tuning. PLoS computational biology, 12(11):e1005175, 2016.

51. Jonathan A Michaels, Stefan Schaffelhofer, Andres Agudelo-Toro, and Hansjōrg Scherberger. A neural network model of flexible grasp movement generation. biorxiv, 2019.

52. Mark M Churchland, John P Cunningham, Matthew T Kaufman, Justin D Foster, Paul Nuyujukian, Stephen I Ryu, and Krishna V Shenoy. Neural population dynamics during reaching. Nature, 487(7405):51–56, 2012.

53. Byron M. Yu, John P. Cunningham, Gopal Santhanam, Stephen I. Ryu, Krishna V. Shenoy, and Maneesh Sahani. Gaussian-Process Factor Analysis for Low-Dimensional Single-Trial Analysis of Neural Population Activity. Journal of Neurophysiology, 102(1):614–635, July 2009.

54. Joshua B Tenenbaum, Vin De Silva, and John C Langford. A global geometric framework for nonlinear dimensionality reduction. science, 290(5500):2319–2323, 2000.

55. Ege Altan, Sara A Solla, Lee E Miller, and Eric J Perreault. Estimating the dimensionality of the manifold underlying multi-electrode neural recordings. PLoS computational biology, 17(11):e1008591, 2021.

56. Stefano Recanatesi, Gabriel Koch Ocker, Michael A Buice, and Eric Shea-Brown. Dimensionality in recurrent spiking networks: Global trends in activity and local origins in connectivity. PLoS computational biology, 15(7):e1006446, 2019.

57. John L Horn. A rationale and test for the number of factors in factor analysis. Psychometrika, 30(2):179–185, 1965.

58. Gopal Santhanam, Byron M Yu, Vikash Gilja, Stephen I Ryu, Afsheen Afshar, Maneesh Sahani, and Krishna V Shenoy. Factor-analysis methods for higher-performance neural prostheses. Journal of neurophysiology, 102(2):1315– 1330, 2009.

59. Matthias Scholz, Martin Fraunholz, and Joachim Selbig. Nonlinear principal component analysis: neural network models and applications. In Principal manifolds for data visualization and dimension reduction, pages 44–67. Springer, 2008.

60. Peter Dunn-Rankin. The similarity of lower-case letters of the english alphabet. Journal of Verbal Learning and Verbal Behavior, 7(6):990–995, 1968.

61. Etay Hay, Albert Gidon, Michael London, and Idan Segev. A theoretical view of the neuron as an input–output computing device. Dendrites, pages 439–464, 2016.

62. Greg Stuart, Nelson Spruston, and Michael Hāusser. Dendrites. Oxford University Press, 03 2016.

63. Joshua T Dudman and Charles R Gerfen. The basal ganglia. In The rat nervous system, pages 391–440. Elsevier, 2015.

64. Aryn H Gittis, Bryan M Hooks, and Charles R Gerfen. Basal ganglia circuits. In Neural Circuit and Cognitive Development, pages 221–242. Elsevier, 2020.

65. Adam G. Carter and Bernardo L. Sabatini. State-Dependent Calcium Signaling in Dendritic Spines of Striatal Medium Spiny Neurons. Neuron, 44(3):483–493, October 2004.

66. Matthew A. Smith and Adam Kohn. Spatial and Temporal Scales of Neuronal Correlation in Primary Visual Cortex. Journal of Neuroscience, 28(48):12591–12603, 2008.

67. Daniel A. Dombeck, Michael S. Graziano, and David W. Tank. Functional Clustering of Neurons in Motor Cortex Determined by Cellular Resolution Imaging in Awake Behaving Mice. Journal of Neuroscience, 29(44):13751–13760, November 2009.

68. Andreas Klaus, Gabriela J. Martins, Vitor B. Paixao, Pengcheng Zhou, Liam Paninski, and Rui M. Costa. The Spatiotemporal Organization of the Striatum Encodes Action Space. Neuron, 95(5):1171–1180.e7, 2017.

69. Saurabh Vyas, Matthew D Golub, David Sussillo, and Krishna V Shenoy. Computation through neural population dynamics. Annual Review of Neuroscience, 43:249, 2020.

70. David L Barack and John W Krakauer. Two views on the cognitive brain. Nature Reviews Neuroscience, 22(6):359– 371, 2021.

71. Elad Ganmor, Ronen Segev, and Elad Schneidman. A thesaurus for a neural population code. Elife, 4:e06134, 2015.

72. Yuke Yan, James M Goodman, Dalton D Moore, Sara A Solla, and Sliman J Bensmaia. Unexpected complexity of everyday manual behaviors. Nature communications, 11(1):1–8, 2020.

73. Trygve E. Bakken, Nikolas L. Jorstad, Qiwen Hu, Blue B. Lake, Wei Tian, Brian E. Kalmbach, Megan Crow, Rebecca D. Hodge, Fenna M. Krienen, Staci A. Sorensen, Jeroen Eggermont, Zizhen Yao, Brian D. Aevermann, Andrew I. Aldridge, Anna Bartlett, Darren Bertagnolli, Tamara Casper, Rosa G. Castanon, Kirsten Crichton, Tanya L. Daigle, Rachel Dalley, Nick Dee, Nikolai Dembrow, Dinh Diep, Song-Lin Ding, Weixiu Dong, Rongxin Fang, Stephan Fischer, Melissa Goldman, Jeff Goldy, Lucas T. Graybuck, Brian R. Herb, Xiaomeng Hou, Jayaram Kancherla, Matthew Kroll, Kanan Lathia, Baldur van Lew, Yang Eric Li, Christine S. Liu, Hanqing Liu, Jacinta D. Lucero, Anup Mahurkar, Delissa McMillen, Jeremy A. Miller, Marmar Moussa, Joseph R. Nery, Philip R. Nicovich, Sheng-Yong Niu, Joshua Orvis, Julia K. Osteen, Scott Owen, Carter R. Palmer, Thanh Pham, Nongluk Plongthongkum, Olivier Poirion, Nora M. Reed, Christine Rimorin, Angeline Rivkin, William J. Romanow, Adriana E. Sedeño-Cortés, Kimberly Siletti, Saroja Somasundaram, Josef Sulc, Michael Tieu, Amy Torkelson, Herman Tung, Xinxin Wang, Fangming Xie, Anna Marie Yanny, Renee Zhang, Seth A. Ament, M. Margarita Behrens, Hector Corrada Bravo, Jerold Chun, Alexander Dobin, Jesse Gillis, Ronna Hertzano, Patrick R. Hof, Thomas Hōllt, Gregory D. Horwitz, C. Dirk Keene, Peter V. Kharchenko, Andrew L. Ko, Boudewijn P. Lelieveldt, Chongyuan Luo, Eran A. Mukamel, António Pinto-Duarte, Sebastian Preissl, Aviv Regev, Bing Ren, Richard H. Scheuermann, Kimberly Smith, William J. Spain, Owen R. White, Christof Koch, Michael Hawrylycz, Bosiljka Tasic, Evan Z. Macosko, Steven A. McCarroll, Jonathan T. Ting, Hongkui Zeng, Kun Zhang, Guoping Feng, Joseph R. Ecker, Sten Linnarsson, and Ed S. Lein. Comparative cellular analysis of motor cortex in human, marmoset and mouse. Nature, 598:111–119, 2021.

74. Stephen H. Scott and John F. Kalaska. Reaching Movements With Similar Hand Paths But Different Arm Orientations. I. Activity of Individual Cells in Motor Cortex. Journal of Neurophysiology, 77:826–852, February 1997.

75. Anil Cherian, Max O Krucoff, and Lee E Miller. Motor cortical prediction of emg: evidence that a kinetic brainmachine interface may be robust across altered movement dynamics. Journal of neurophysiology, 106(2):564–575, 2011.

76. Benjamin Y Hayden, Hyun Soo Park, and Jan Zimmermann. Automated pose estimation in primates. American journal of primatology, page e23348, 2021.

77. Emily Jane Dennis, Ahmed El Hady, Angie Michaiel, Ann Clemens, Dougal R Gowan Tervo, Jakob Voigts, and Sandeep Robert Datta. Systems neuroscience of natural behaviors in rodents. Journal of Neuroscience, 41(5):911–919, 2021.

78. M Franch, S Yellapantula, A Parajuli, N Kharas, A Wright, B Aazhang, and V Dragoi. Visuo-frontal interactions during social learning in freely moving macaques. Nature, pages 1–8, 2024.

79. William Bialek. On the dimensionality of behavior. Proceedings of the National Academy of Sciences, 119(18):e2021860119, 2022.

80. P. W. Anderson. More Is Different. Science, 177(4047):393–396, August 1972. Publisher: American Association for the Advancement of Science.

81. Paul Humphreys. Emergence: A philosophical account. Oxford University Press, 2016.

82. Junchol Park, Luke T. Coddington, and Joshua T. Dudman. Basal Ganglia Circuits for Action Specification. Annual Review of Neuroscience, 43(1):485–507, July 2020.

83. Suzanne N. Haber. The primate basal ganglia: parallel and integrative networks. Journal of Chemical Neuroanatomy, 26(4):317–330, 2003.

84. Akash Umakantha, Rudina Morina, Benjamin R Cowley, Adam C Snyder, Matthew A Smith, and M Yu Byron. Bridging neuronal correlations and dimensionality reduction. Neuron, 109(17):2740–2754, 2021.

85. Mehrdad Jazayeri and Srdjan Ostojic. Interpreting neural computations by examining intrinsic and embedding dimensionality of neural activity. Current opinion in neurobiology, 70:113–120, 2021.

86. David Dahmen, Stefano Recanatesi, Gabriel K Ocker, Xiaoxuan Jia, Moritz Helias, and Eric Shea-Brown. Strong coupling and local control of dimensionality across brain areas. Biorxiv, pages 2020–11, 2020.

87. Erik Hermansen, David A. Klindt, and Benjamin A. Dunn. Uncovering 2-d toroidal representations in grid cell ensemble activity during 1-d behavior. bioRxiv, 2022.

88. Timothy EJ Behrens, Timothy H Muller, James CR Whittington, Shirley Mark, Alon B Baram, Kimberly L Stachenfeld, and Zeb Kurth-Nelson. What is a cognitive map? organizing knowledge for flexible behavior. Neuron, 100(2):490–509, 2018.

89. Edvard I. Moser, Emilio Kropff, and May-Britt Moser. Place Cells, Grid Cells, and the Brain’s Spatial Representation System. Annual Review of Neuroscience, 31(1):69–89, 2008.

90. Wei Guo, Jie J Zhang, Jonathan P Newman, and Matthew A Wilson. Latent learning drives sleep-dependent plasticity in distinct ca1 subpopulations. bioRxiv, 2020.

91. Juan A Gallego, Tamar R Makin, and Samuel D McDougle. Going beyond primary motor cortex to improve brain–computer interfaces. Trends in neurosciences, 2022.

92. Chethan Pandarinath and Sliman J Bensmaia. The science and engineering behind sensitized brain-controlled bionic hands. Physiological Reviews, 102(2):551–604, 2022.

93. Cecilia Gallego-Carracedo, Matthew G Perich, Raeed H Chowdhury, Lee E Miller, and Juan Á lvaro Gallego. Local field potentials reflect cortical population dynamics in a region-specific and frequency-dependent manner. Elife, 11:e73155, 2022.

94. Junchol Park, James W Phillips, Jian-Zhong Guo, Kathleen A Martin, Adam W Hantman, and Joshua T Dudman. Motor cortical output for skilled forelimb movement is selectively distributed across projection neuron classes. Science advances, 8(10):eabj5167, 2022.

95. Mayank Kabra, Alice A. Robie, Marta Rivera-Alba, Steven Branson, and Kristin Branson. JAABA: interactive machine learning for automatic annotation of animal behavior. Nature Methods, 10(1):64–67, January 2013. Number: 1 Publisher: Nature Publishing Group.

96. Jason E Osborne and Joshua T Dudman. Rivets: a mechanical system for in vivo and in vitro electrophysiology and imaging. PloS one, 9(2):e89007, 2014.

97. F Pedregosa, G Varoquaux, A Gramfort, V Michel, B Thirion, O Grisel, M Blondel, P Prettenhofer, R Weiss, V Dubourg, et al. “ scikit-learn: Machine learning in python,” journal of machine learning research, vol. 12, p. 2011.

98. Adam Paszke, Sam Gross, Soumith Chintala, Gregory Chanan, Edward Yang, Zachary DeVito, Zeming Lin, Alban Desmaison, Luca Antiga, and Adam Lerer. Automatic differentiation in pytorch. In NIPS-W, 2017.

99. Gamaleldin F. Elsayed, Antonio H. Lara, Matthew T. Kaufman, Mark M. Churchland, and John P. Cunningham. Reorganization between preparatory and movement population responses in motor cortex. Nature Communications, 7(1):13239, December 2016.

100. Joshua I Glaser, Ari S Benjamin, Raeed H Chowdhury, Matthew G Perich, Lee E Miller, and Konrad P Kording. Machine learning for neural decoding. Eneuro, 7(4), 2020.

101. Francis R Bach and Michael I Jordan. Kernel independent component analysis. Journal of machine learning research, 3(Jul):1–48, 2002.

